# Cholesterol is a strong promotor of an α-Synuclein membrane binding mode that accelerates oligomerization

**DOI:** 10.1101/725762

**Authors:** Martin Jakubec, Espen Bariås, Samuel Furse, Morten L. Govasli, Vinnit George, Diana Turcu, Igor Iashchishyn, Ludmilla Morozova-Roche, Øyvind Halskau

## Abstract

Dysregulation of the biosynthesis of cholesterol and other lipids has been implicated in neurological diseases, including Parkinson's disease, where the misfolding of membrane-associated α-Synuclein is a key molecular event. Recent research also suggests that α-Synuclein aggregation is influenced by the lipid environment. The exact molecular mechanisms responsible for cholesterol’s effect on α-Synuclein binding to lipids and how this binding may affect α-Synuclein oligomerization and fibrillation remain elusive, as does the relative importance of cholesterol versus other lipid factors. We probed the interactions and fibrillation behaviour of α-Synuclein using SMA nanodiscs, containing zwitterionic and anionic lipid model systems with and without cholesterol. SPR and ThT fluorescence assays were then employed to monitor α-Synuclein binding, as well as fibrillation in the absence and presence of membrane models. ^1^H-^15^N correlated NMR was used to monitor the fold of α-Synuclein in response to nanodisc binding, and we determined individual residue apparent affinities for the nanodisc-contained bilayers. Cholesterol inhibited α-Synuclein interaction with lipid bilayers. We also find that cholesterol significantly promotes α-Synuclein fibrillation, with a more than 20-fold reduction of lag-times before fibrillation onset. When α-Synuclein-bilayer interactions were analysed for individual residues by solution-state NMR, we observed two different effects of cholesterol. In nanodiscs made of DOPC, cholesterol modulated the NAC part of α-Synuclein, leading to stronger interaction of this region with the lipid bilayer. In contrast, in the nanodiscs comprising DOPC, DOPE and DOPG, the NAC part was mostly unaffected by cholesterol, while the binding of the N-terminal and the C-terminal were both inhibited.

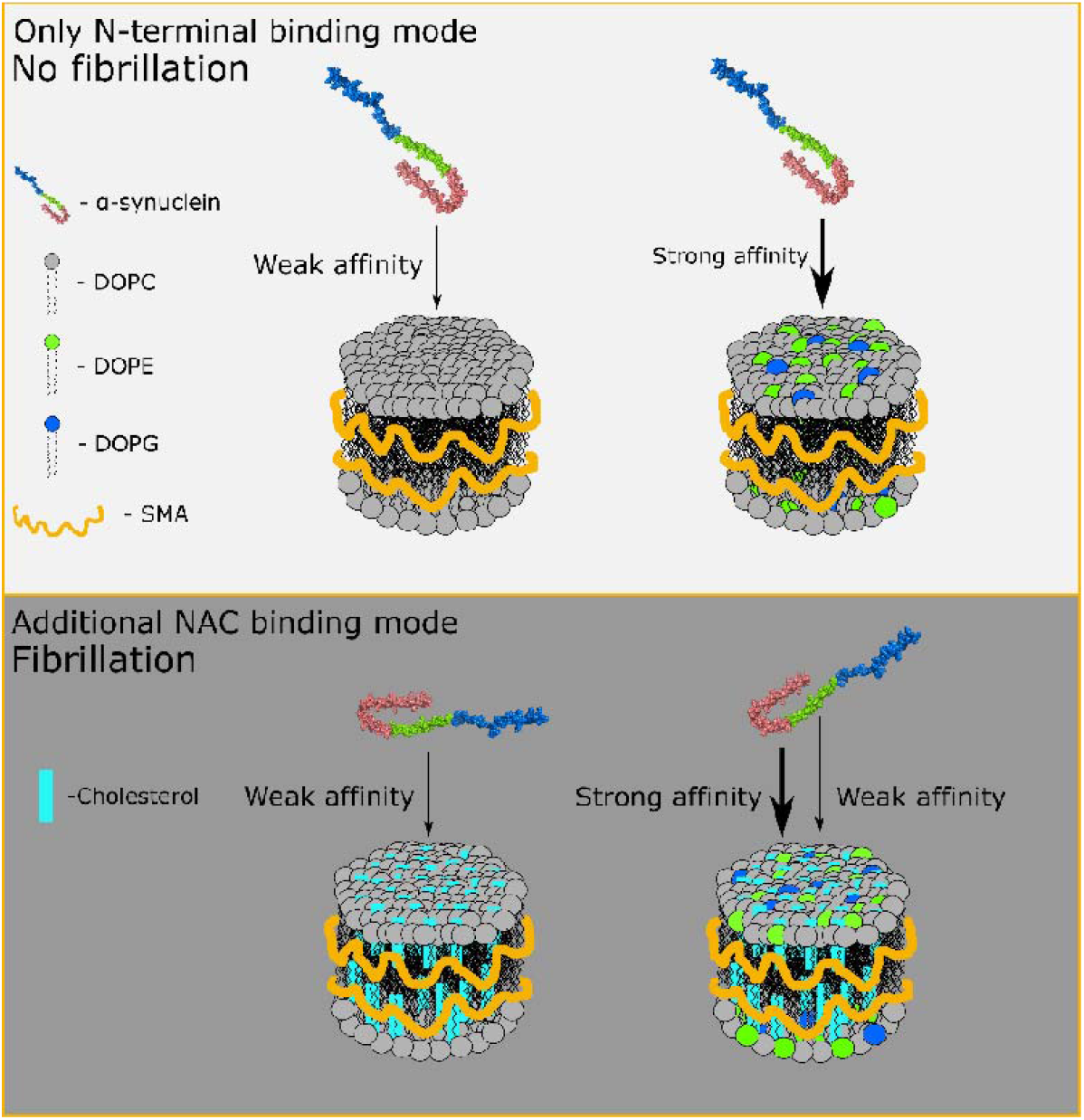

## Introduction

Parkinson’s disease (PD) is a protein misfolding disease associated with a conversion of α-Syn from its soluble state into fibrillar aggregates. During this conversion, α-Syn forms soluble, toxic oligomers that are implicated in the dopaminergic neuron cell death associated with clinical manifestations of PD ^1^. The exact function of native α-Syn is not fully understood, nor are the driving forces governing its misfolding, oligomerization and fibrillation, including the specific roles of individual residues within the protein. It has been proposed that α-Syn plays a role in synaptic plasticity allowing ‘kiss-and-run’ neurotransmitter release, where the protein assists vesicles containing the neurotransmitters to fuse with the synaptic membrane transiently so that it can deliver its cargo before disengaging^2^. Unfortunately, the role of the interaction between α-Syn and lipid bilayers is unclear as a consistent pattern of behaviour has not been observed (review ^3^). Research, including a recent comprehensive work by Viennet *et al* ^4^, indicate that two important membrane general characteristics are affecting the interaction between α-Syn and lipid membrane; its net charge and physical state. A higher abundance of anionic lipids, like phosphatidylserine (PS) or phosphatidylglycerol (PG), leads to a higher affinity of α-Syn and a slower aggregation ^4,5^. Likewise, pronounced membrane curvature and smaller lipid head group size seem to promote the interaction of α-Syn ^4,6,7^. With this is in mind, there is a notable lack of knowledge regarding α-Syn interaction with cholesterol, the most abundant mammalian component of the plasma membrane (PM).

Cholesterol constitutes about 50% of the mass of PMs in mammals and has a considerable effect on its physical properties, including its fluidity and permeability ^8,9^. When residing in the bilayer, cholesterol is only marginally accessible to the solvent or peripheral protein interaction as it is primarily its −OH group that is exposed ^10^. However, cholesterol has prominent effects on how other lipids are organized, and may in this way confer larger effects than its small exposure would initially suggest. Cholesterol is particularly abundant in the brain and its dysregulation has been linked to several neurodegenerative diseases, including AD ^11,12^, Huntington’s disease ^13^ or Niemann-Pick disease type C ^14,15^. Recent evidence suggests that cholesterol is more abundant in the inner monolayer of the synaptic membrane, where α-Syn makes contact with the PM as part of its function ^16^. It has also been observed that the cholesterol abundance in the PM of neurons decreases with age, which in turn could drive or confer vulnerability towards neurodegeneration ^17^. This age-dependent loss of cholesterol from the PM seems to affect the release of neurotransmitters by hindering the fusion of presynaptic vesicles ^18^. However, the clinical connection between PD and cholesterol abundance and distribution remains unclear. There is evidence that higher level of serum cholesterol is associated with a higher risk of PD ^19^, a decrease in the risk of PD ^20^, or is not associated with PD at all ^3,21^.

There are also contradictory results regarding the interaction of α-Syn and cholesterol at the molecular level. It has been reported that there is a cholesterol binding site in the Non-Amyloid-Component (NAC) of α-Syn and that cholesterol is essential for the formation of ‘amyloid pores’ ^22–24^, which is relatable to the toxic effects that has been proposed for α-Syn oligomers. However, recent work has shown that the presence of cholesterol inhibits the binding of α-Syn to membranes comprising either zwitterionic or anionic lipids ^25,26^. The apparent discrepancy in affinity between α-Syn and cholesterol, and the interaction of α-Syn and lipid head groups has, to our knowledge, not been explained adequately. Despite contradictory and incomplete evidence, current knowledge strongly suggests a role for cholesterol in both the normal function and misfolding of α-Syn. We hypothesise that there is an interplay between cholesterol abundance of the membrane and the reversible and dynamic α-Syn binding, the protein conformational response, and ultimately the protein oligomerization and fibrillation outcome. We further propose that these changes in the protein conformation are primary driven by interaction between the NAC region and cholesterol.

The investigation of reversible protein-membrane interactions often relies on lipid vesicles or supported lipid bilayers ^27^. However, such lipid assemblies limits the interpretation and application of many methods including solution-state NMR, steady-state affinity measurements and fibrillation studies ^28^. Nanodiscs, consisting of circular patches of lipid bilayer surrounded by a polymer belt, have emerged as an alternative to vesicles and have been successfully used in structural studies of membrane proteins and anchored peripheral proteins ^28,29^. As they can be prepared with narrow size distributions, well-defined lipid compositions, and are stable in solution, they are promising tools for studying reversible protein-lipid interactions ^29,30^. In this study, we use styrene-maleic acid (SMA) nanodiscs as the primary membrane model system for our studies. These nanodiscs have properties similar to those that are prepared using a protein belt to scaffold the lipids, but allow a detergent-free sequestration of lipids directly from vesicle or even native membranes ^29^. Recently, protein-scaffolded nanodiscs were used to investigate the role of charge and fluidity on α-Syn binding and aggregation ^4^. The study notes that while high-resolution solution state NMR is made possible by nanodiscs and yields sequence-specific binding information, it does not allow direct detection of α-Syn’s membrane-bound state ^4^. However, their comprehensive work allowed them to propose a model for α-Syn membrane interaction and its effect on fibrillation that focused on the protein competing for favourable binding sites on fluid patches of net negative charge. In their model, increased fibrillation occurred when α-Syn competed for the most favourable binding sites, bringing exposed NAC regions together ^4^. Given the importance of cholesterol for α-Syn binding, and the implication of the NAC region, further detailed investigation focusing on the effect of cholesterol is warranted.

Here we present a comparative lipid-dependent binding and fibrillation study of recombinant α-Syn using SMA lipid nanodiscs consisting of zwitterionic DOPC and a membrane model containing DOPC, PE and PG in a 4:3:1 ratio. Each model was also prepared with 30% (w/w) cholesterol. We have used 2D ^1^H/^15^N-HSQC NMR to monitor α-Syn-membrane binding of individual amino acid by titrating lipid nanodiscs into samples of ^15^N labelled α-Syn. We have, where possible, used the increasing width of the resonance base peak of detectable amino-acids with an increasing amount of lipid nanodiscs in the system, as an indicator for how the presence of nanodiscs affect α-Syn. This allowed us to estimate residue-specific apparent binding constants for ~40% of residues evenly distributed within the α-Syn sequence, using a formalism adapted from Shortridge *et al.* ^31^. These binding constants were compared with steady state affinity measurements obtained using surface plasmon resonance (SPR) with lipid vesicles 100 nm across. Lastly, we monitored the effect of cholesterol in lipid nanodiscs on oligomerization rates of α-Syn by using ThT and a novel tetraphenylethene tethered with triphenylphosphonium (TPE-TPP) fluorescence marker ^32^, potentially allowing earlier detection of the onset of fibrillation than the standard ThT approach.

Cholesterol strongly promotes α-Syn fibrillation for both lipid models investigated here, suggesting that the cholesterol-mediated binding mode seeds this process. The cholesterol effect on both overall protein affinity and individual residue in the N-terminal and NAC region is dependent on the exact composition of the model membranes investigated. Our results indicate that planar, zwitterionic and cholesterol rich sites, are potentially important sites for fibrillation.

## Results

### Binding of α-Syn monomers to lipid vesicles is inhibited by cholesterol

Net negative bilayer charge has been established as a primary determinant for reversible protein-membrane interactions in general ^33–36^, and also for α-Syn ^4,7,37^. We therefore prepared vesicles representing either fluid zwitterionic or fluid anionic bilayers. The zwitterionic vesicles were prepared using DOPC alone, while the anionic model was more complex using DOPC:PE:PG in a 4:3:1 molar ratio (structure of lipids shown in <u>Figure S1A</u>). The latter model confers net charge at most pHs and has a PE content similar to that of PM membrane. Each model was also prepared in the presence or absence of 30% (w/w) cholesterol, yielding a total of four different lipid compositions. The steady-state affinity of monomeric (>95% according to SEC, <u>Figure S2A</u>) α-Syn for the lipid vesicles was then investigated using a Biacore T200 instrument in multiple cycle experiment runs.

A representative SPR plot is shown in Figure 1A-D. The dissociation equilibrium constant, K_D_, of α-Syn was estimated using Equation 1 (Figure 1D). The dissociation constant for DOPC mixtures was approximately 7.2 times higher (1072 ± 137 nM) than for DOPC:PE:PG vesicles (148 ± 36 nM). This is consistent with earlier studies that found that α-Syn has a relatively low affinity for the DOPC bilayer, while the presence of anionic PG and zwitterionic PE increased the affinity of α-Syn to the lipid vesicles ^37–40^. Vesicles containing 30% (w/w) cholesterol showed an almost a two-fold increase in the K_D_ (2138 ± 322 nM for DOPC and 273 ± 36 nM for DOPC:PE:PG vesicles). These values indicate that cholesterol has a general inhibitory effect as the K_D_ was increased in both lipid models. This led us to a question whether the presence of cholesterol affects α-Syn misfolding and aggregation behaviour. However, quasi-stable vesicle models are not suitable for fibrillation studies that often span days ^41^. Therefore, we explored this question using lipid nanodiscs.

**Figure 1:**
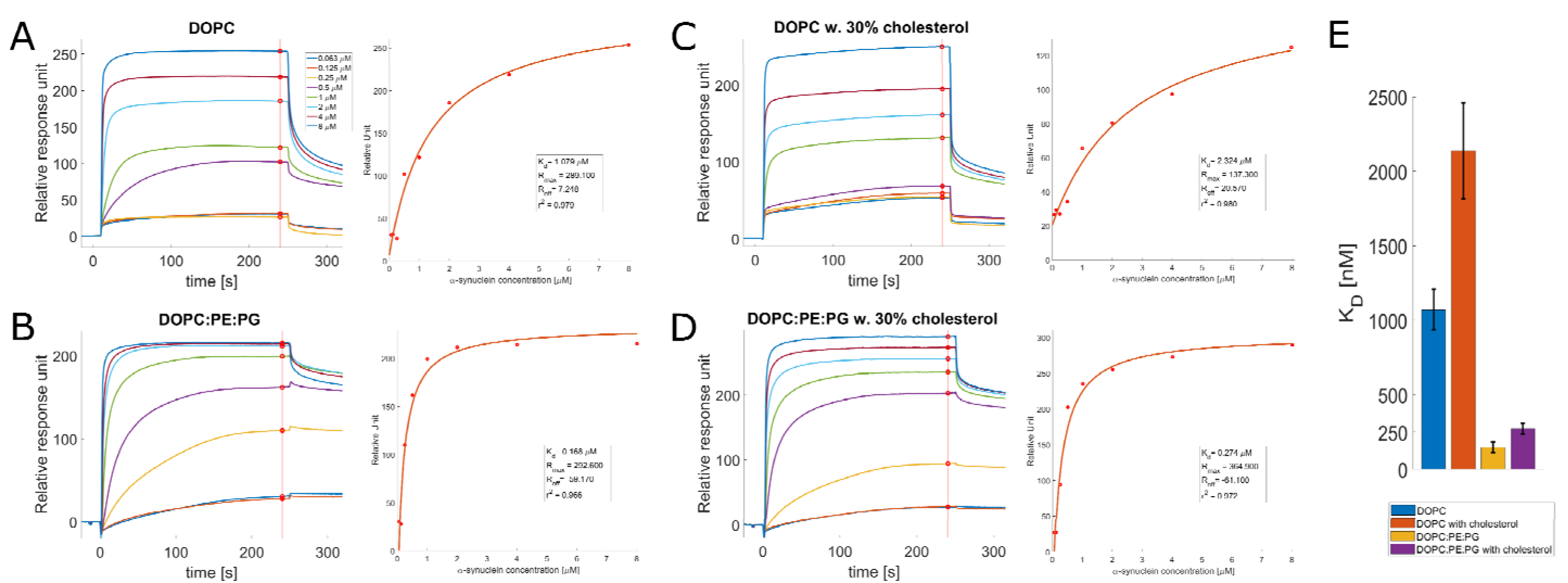
Monomeric α-Syn binding to vesicles is inhibited by cholesterol. Representative SPR plots of α-Syn interaction with immobilized lipid layers of DOPC (A) and mixtures of DOPC:PE:PG (4:3:1) (B), without and with the addition of 30% (w/w) cholesterol (C,D, respectively). Equilibrium values are indicated with red circles where the vertical lines intersect the sensorgrams. Red line represent fitting into Equation 1. (E) Mean K_D_ values for each lipid group. Each value represents 3 to 5 replicates and error bars depict standard deviations.

### The presence of cholesterol in the lipid nanodiscs influences both lag time and rate of fibrillation of α-Syn

It has been reported that cholesterol can promote interaction of α-Syn oligomers and zwitterionic PC and PE lipids prepared as vesicles and sonicated lipid dispersions ^42,43^. However, the effect of cholesterol on α-Syn aggregation is not known for lipid model systems with planar bilayers. Moreover, previous studies used either lipid vesicles, which are not suitable for long (more than 24 hours) aggregation experiments, or seeding, which will lead to faster aggregation rates, but also increase batch-to-batch variability. We therefore prepared nanodiscs from 100 nm vesicles of the same compositions as used for the SPR affinity measures. This was done by adding SMA to 1% w/v, and overnight incubation followed by size exclusion chromatography (SEC) purification. Although the presence of cholesterol and PE in the vesicles inhibited proper nanodisc formation, we were able to optimize the process as described in Supplementary Information. Nanodisc elution times and dimensions as determined by SEC and DLS, respectively, are summarized in Table S1. Inclusion of cholesterol into the nanodiscs where verified using LC MS/MS (Figure S3).

We then were able to test how the presence of cholesterol embedded in nanodiscs affected the fibrillation rate of α-Syn. ThT and TPE-TPP fluorescence assays were used to measure the fibrillation of α-Syn both in the absence and presence of lipid nanodiscs. While ThT is a fluorescence probe that is widely used to characterize the aggregation of various amyloidogenic peptides, also in the presence of lipid bilayers ^44–46^. TPE-TPP, is to our knowledge untested as a fluorescence marker in the presence of lipids. The compound is a structural ThT analogue developed by Leung *et al.* ^32^. It binds to fibrils by a similar mechanism as ThT, but it is two orders of magnitude more sensitive and therefore potentially an earlier reporter of fibrillation ^47^.

We adopted a high-throughput technique, using 384 well plates and incubation time of up to 150 hours ^48^. First, we evaluated the experiment by cumulative curve plot ^48^. Briefly, each well was annotated as either positive or negative, based on its fluorescence signal. A well is considered positive when its fluorescence signal exceeds three times the median value of the control wells (without α-Syn). The percentage of positive wells against time is then plotted (Figure 2A), and from this the lag time for each experimental condition can be read directly. The lag time of α-Syn, 20 hours, increased in the presence of DOPC and DOPC:PE:PG lipid nanodiscs to 50 and 60 hours, respectively. However, with cholesterol present in the lipid nanodiscs, the lag time dropped to 3 and 2 hours, respectively. We then evaluated the experiment more in-depth, by fitting the fluorescence signals from each well independently to a two-step Finke-Watzky model ^49^ (Equation 2). From this model, we obtained the t_N_ (lag time, Equation 3) and ν (fibrillation rate). The ratio of t_N_/t_N_-control and ν/ν-control is presented in Figure 2B and C.

**Figure 2:**
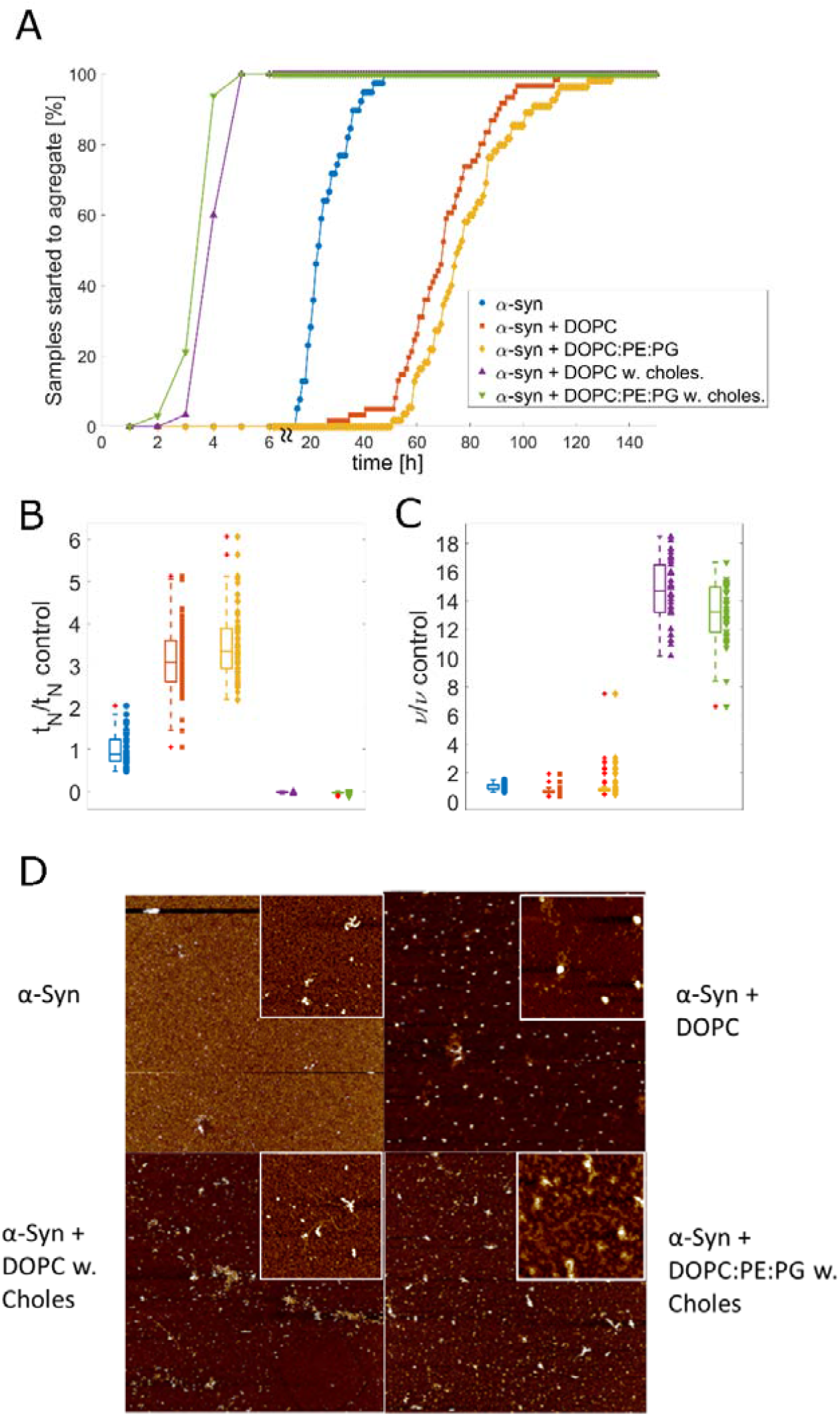
Cholesterol promotes early oligomerization. (A) The cumulative distribution of ThT positive wells (total of 30-61 repeats from 3 independent experiments). (B) Representative examples of single oligomerization curves for each sample, with fits calculated based on Equation 2 and 3. (C) Ratio of t_N_/t_N control_, which represent lag time ratio (relative primary nucleation rate). (C) Ratio of ν/ν_control_, which represents the relative growth rate. (B, C) Central mark of the box indicates median, the top and bottom of the box indicates 25th and 75th percentiles, respectively. The whiskers extend to extreme data points, outliers are indicated by red ‘+’ symbol. (D) Representative AFM pictures of α-Syn in the presence and absence of nanodiscs with and without cholesterol. The lipid composition of the nanodiscs used in each case is indicated next to the figure panels. The images are all acquired at 256px · 256px resolution during the lag-phases of the ThT-monitored fibrillation assays. Each panel have a 10 μm focus, while the zoom insets all have a focus of 2 μm. Vertical scales span, from darkest to brightest colour: α-Syn (2.6 nm, inset 4.0 nm), α-Syn + DOPC (16.5 nm, inset 19.8 nm), α-Syn DOPC with Cholesterol (6.6 nm, inset 3.4 nm), α-Syn + DOPC:PE:PG with Cholesterol (7.5 nm, inset 8.8 nm).

Using the TPE-TPP fluorescence dye, we were able to detect the early stages of fibrillation of α-Syn alone (<u>Figure S2B</u>). However, with the addition of lipid nanodiscs we observed a loss of fluorescence (<u>Figure S4</u>). This suggests the release of TPE-TPP from the cross-β sheets of α-Syn and its interaction with the lipid nanodiscs. TPE-TPP was therefore deemed unsuitable for fibrillation studies in the presence of our model system of choice but did serve well as an early, sensitive marker for fibrillation events. The results presented below are therefore derived from ThT-based assays only.

We also monitored the morphology of α-Syn during the lag phase using atomic force microscopy (Figure 2D). For α-Syn alone we could not observe any fibrils, as α-Syn was observed only as a granular background using a focus of both 10 μm and 2 μm (Figure 2D insets). From α-Syn + DOPC, we can clearly observe the DOPC-nanodisc as spheres of 50-100 nm, larger than the diameters in Table S1. The background is otherwise relatively clear when comparing to α-Syn alone. In the 2 μm focus, however, we observe what we interpret as some aggregation of α-Syn around the nanodiscs. The same situation, but also with cholesterol present, shows a more diverse morphology, where nanodiscs and α-Syn co-aggregate to a larger extent. The 2 μm focus inset shows an α-Syn fibril. Also in the case of the negatively charged DOPC:PE:PG nanodiscs with cholesterol, we observed a mix of morphologies consistent of discs with associated α-Syn aggregates. The 2 μm inset shows fibril morphologies, albeit shorted and more diffuse than in the case of DOPC with cholesterol.

The t_N_ ratio (lag time ratio) is three times higher when nanodiscs (without cholesterol) are present during fibrillation (3.10 ± 0.77 for α-Syn and DOPC, 3.53 ± 0.84 for α-Syn and DOPC:PE:PG and 1 ± 0.38 for α-Syn alone). However, the fibrillation rate ratio, ν, did not change significantly (1 ± 0.22 for α-Syn alone, 0.71 ± 0.22 for α-Syn and DOPC, and 1.13 ±1.05 for α-Syn and DOPC:PE:PG, t-test P< 0.01). When cholesterol was present the t_N_ ratio approached zero as the fluorescence started to increase almost immediately after the addition of lipid nanodiscs (−0.01 ± 0.20 for α-Syn and DOPC with 30% cholesterol and −0.03 ± 0.03 for α-Syn and DOPC:PE:PG with 30% cholesterol). We also observed a significant increase of the ν ratio to approximately 14x in the presence of cholesterol compared to the control (14.59 ± 2.26 for α-Syn and DOPC with 30% cholesterol and 13.11 ± 2.13 for α-Syn and DOPC:PE:PG with 30% cholesterol).

While cholesterol actively promotes fibrillation, the SPR experiment indicated that interaction between vesicles and α-Syn is diminished in the presence of cholesterol (Figure 1). This suggests to us that cholesterol is having an effect similar to seeding; it is either promoting an α-Syn fold which is more prone to aggregation or is stabilising oligomers which are on-pathway to fibrils. This is consistent with the observation that atomic form microscopy did observe aggregation in direct contact with cholesterol-containing nanodiscs (Figure 2D). The presence of cholesterol may also modulate which parts of α-Syn come into contact with and embed in the bilayer. We explored these possibilities further using solution state NMR.

### Cholesterol affects the interaction of α-Syn and lipid bilayer differently in DOPC and DOPC:PE:PG lipid models

To further explore the mechanism of α-Syn and cholesterol interaction we proceeded with NMR studies to determine which amino acids are affected in the presence of lipid nanodiscs. Lipid nanodiscs of the compositions described above were titrated into ^15^N labelled α-Syn and HSQC fingerprints were acquired to monitor changes in the chemical environment of individual residues. Cross-peak assignments were adapted from BMRB entry 6968 ^50^ and 25227 ^51^ and verified using C_α_ and C_β_ chemical shifts from standard triple resonance experiments performed on a double-labelled α-Syn sample. In agreement with other studies^52–54^, we did not observe any chemical shifts of cross-peaks, which would represent a direct observation of changes in the chemical environment. Instead, we observed line (peak) broadening which grows beyond detection in the high concentrations of lipid nanodiscs (Figure 3A). These undetectable peaks, sometimes called invisible states, are ascribed to chemical exchange broadening or efficient relaxation pathways related to molecular motions^52^. Free and lipid-associated α-Syn in exchange, intra-molecular interaction in folding, and inter-molecular oligomerization may all cause line broadening and loss of observable states. We evaluated these invisible states as possible binding curves, using logic presented by Shortridge *et al.* ^31^ for estimating protein/ligand affinity. Between 77 (α-Syn and DOPC:PE:PG with 30% cholesterol) and 82 (α-Syn and DOPC) cross-peaks and their individual line-broadening behaviour in response to lipid nanodiscs were observed for each lipid nanodisc model. Using the PINT software ^55^, we deconvoluted peaks for each assigned amino acid and calculated a residue-specific apparent K_D_ (aK_D_) based on Equation 4 ^31^. From 48 (α-Syn and DOPC:PE:PG with 30% cholesterol) to 67 (α-Syn and DOPC) cross-peaks had good correlations with the model (r > 0.90, Figure 3B).

**Figure 3:**
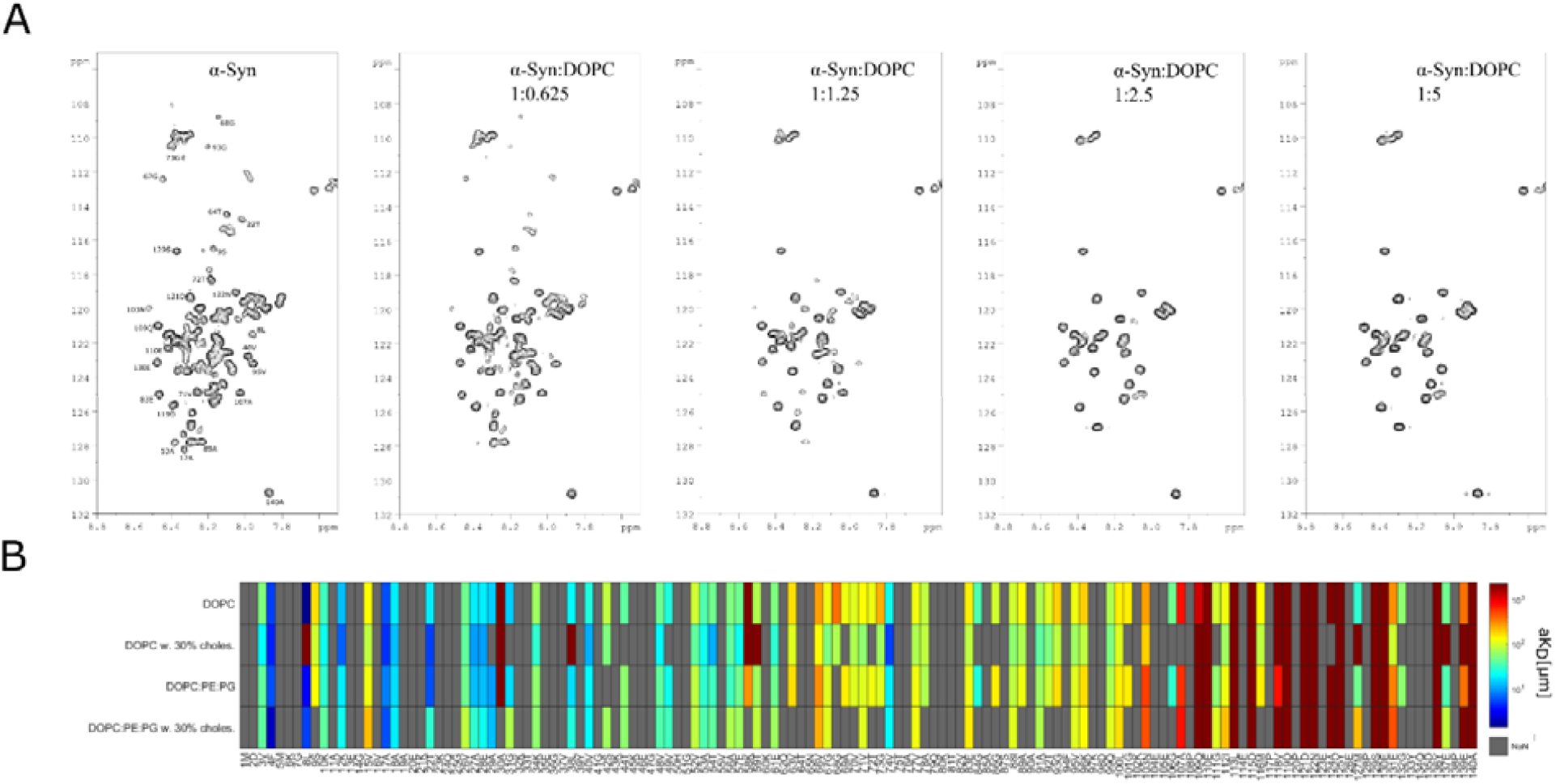
Site specific interaction of α-Syn with lipid nanodiscs with and without cholesterol. (A) HSQC plots of 100 μM α-Syn and DOPC nanodiscs. With increasing lipid nanodisc concentrations (62.5, 125, 250 and 500 μM) from left to right, there is a decrease in a number of observed α-Syn peaks. (B) Heatmap of calculated aK_D_ values for amino acid residues of α-Syn in the presence of lipid nanodiscs from Equation 4. Only the amino acids with r^2^ > 0.90 are shown

In order to visualize the results, each residue-specific aK_D_ was then plotted as a function of lipid nanodisc composition and its position in the α-Syn primary sequence (Figure 3B). High values of aK_D_ (> 500 μM) were observed in the C-terminal part of α-Syn in the presence of all lipid models. The lowest aK_D_ values were observed for N-terminal part of α-Syn suggesting that this part of the molecule is essential for the binding to lipid bilayers. This is consistent with earlier studies ^4,56,57^. Overall, the heat map suggests that there are three distinct regions of α-Syn: the N-terminal with high affinity for lipid membranes, the middle NAC region with an intermediate affinity and the C-terminal with low affinity. This division is expected, as it fits well with a standard description of α-Syn structure ^4,58,59^. Consistent with recent research ^54,60^, most of the observable peaks in the C-terminal region did not change in any way, suggesting most of the C-terminal domain remains unperturbed in a random coil formation. These peaks are depicted in Figure 3B as deep red (See Table S3 for values).

Since cholesterol has such an impact on both binding and fibrillation rates, we wanted to see which parts of α-Syn that were particularly affected by its presence. Changes in the aK_D_ values in the presence of cholesterol was therefore plotted in Figure 4A and 4B and illustrated on the PDB model on Figure 4D. When cholesterol is present in the lipid nanodiscs we could see two different types of interaction behaviour. For the DOPC:PE:PG model we observed a slight increase of aK_D_ in both the amphipathic and acidic region of α-Syn, while the NAC region is mostly unaffected (Figure 4B). The aK_D_ increase was most prominent in the C-terminal region and in several N-terminal region residues. This is in line with what was observed by SPR (Figure 1), where cholesterol seems to inhibit the interaction of α-Syn and the anionic lipid bilayer. In contrast, the trend for DOPC and cholesterol is entirely different (Figure 4A). When cholesterol was present in DOPC lipid nanodiscs we observed a decrease of aK_D_ along almost the whole sequence of α-Syn. This decrease was highest in the NAC region, which suggests that this part was the key α-Syn segment in binding towards DOPC bilayers containing cholesterol. However, the overall increase of aK_D_s for DOPC:PE:PG nanodiscs containing cholesterol was smaller than the overall decrease of aK_D_s in DOPC nanodiscs containing this lipid. This suggests that the effect of including smaller and charged head-groups is higher and more pronounced than the effect of cholesterol on α-Syn binding towards the lipids.

**Figure 4:**
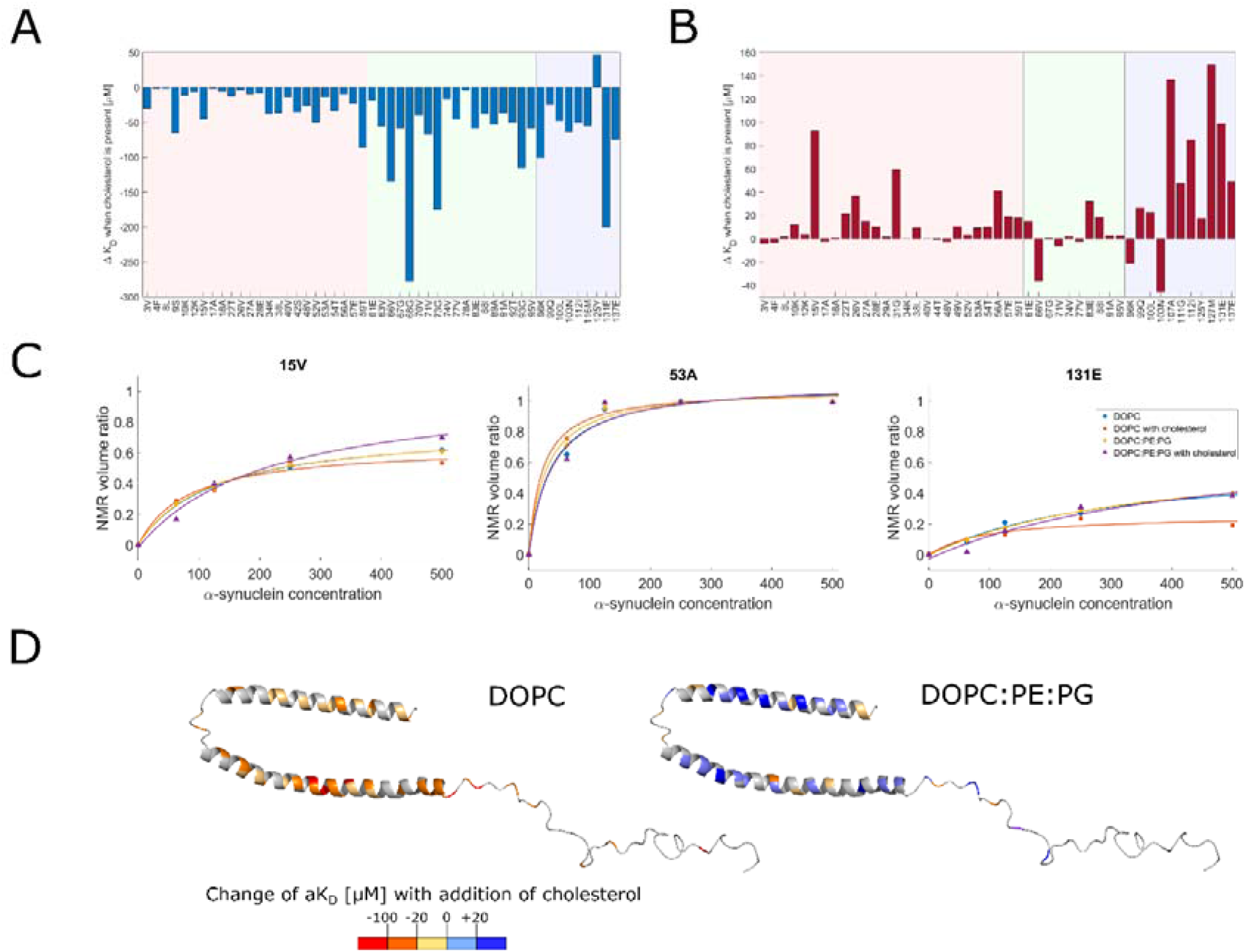
Cholesterol in non-charged nanodiscs promotes site specific α-Syn binding. (A,B) Bar plot of aK_D_ difference for samples with and without cholesterol for DOPC (A) and DOPC:PE:PG (B) lipid nanodiscs. Only amino acids with r^2^ > 0.90 for both samples are shown. The sequence is visually divided into three parts of α-Syn protein amphipathic region, NAC domain and acidic tail (light red, light green, light blue, from left to right). (C) Example of fit for three amino acid fits – V15, A53 and A131 with a medium, high and low aK_D_ (from left to right). Each line represent different lipid environment. (D) Representation of the estimated difference of aK_D_ values after the addition of cholesterol. The representation was made using PDB entry 1XQ8 as a template.

## Discussion

The interest in the role of lipid composition on α-Syn function and dysfunction have prompted research with a variety of membrane model systems ^61^. Although nanodiscs comprising lipids with a variety a charge and acyl-chain saturations have been used in recent α-Syn research ^4,62^, we are unaware of any studies using cholesterol-containing discs. Given the importance of this component of the plasma membrane and its possible role in PD disease at molecular and organism levels, we prepared cholesterol-containing discs using the SMA polymer approach (<u>Figure S1</u>). The presence of cholesterol and PE impaired disc formation, but this was overcome by adjusting the SMA concentration and by having SMA present at vesicle extrusion (more information in Supplementary). SMA discs consequently also proved to be versatile model systems with respect to the lipids that can be included in the bilayer, which allowed us to investigate the effect of cholesterol and other lipids on α-Syn binding and fibrillation. It also allowed us to investigate the protein conformational response using NMR, but the problem with protein states that are invisible to NMR remains due to a combination of complex chemical exchange between protein states and line broadening effects due to efficient relaxation ^4,63^. For fibrillation experiments, the nanodiscs removed reliance on unstable and/or insoluble vesicles, allowing much longer experiments to be pursued. In contrast to MBP-scaffolded nanodiscs, SMA may extract membrane patches directly from live cell membranes and form nanodiscs with a biomolecular profile, including lipids, corresponding to the cell line in use ^29^.

The SPR binding experiments revealed relatively low affinity between monomeric α-Syn and DOPC vesicles. However, a change in composition towards higher net charge and smaller head groups (from pure DOPC to a mixture of DOPC:PE:PG 4:3:1) lead to substantial increase in affinity (Figure 1). The higher affinity between α-Syn and bilayers containing DOPC:PE:PG in 4:3:1 mix can primarily be attributed to the initial electrostatic interaction between the N-terminal part of α-Syn and anionic lipid headgroups ^38,64^. Corresponding bilayers containing cholesterol caused an almost two-fold increase of K_D_ in both lipid models. However, the K_D_ for DOPC:PE:PG with 30% cholesterol was still almost four times lower than for the DOPC membrane model. This suggests that a tighter and more ordered packing of lipids has a negative impact on the α-Syn bilayer interaction, although the presence of net negative charge is a stronger determinant in the current setup. Our findings are consistent with recent studies reporting that cholesterol has negative effect on α-Syn binding ^25^, and with reports finding that bilayer fluidity is an important driver for binding alongside charge ^4,42^. The binding of α-Syn may be further promoted by its potential for desolvating exposed hydrophobic sites between small headgroups the bilayer, alleviating the entropic cost associated with this and at the same time reducing curvature elastic stress ^39,65^. For α-Syn binding to PE-containing lipids these exposed sites may be more important in the absence of strong electrostatic interactions.

The nanodiscs employed in this study made it possible to investigate the effect of membrane components on fibrillation rates. As described above, we modified the high throughput method described by Afitska *et al.* ^48^, where each well in a 384-well plate was counted either as positive or negative with respect to fibrillation. The fraction of positive wells as a function of time was fitted to the Finke-Watzky two-step model (Equation 2) ^49^. This approach avoids experimental sensitivity to initial conditions that are difficult to control for. The onset of α-Syn fibrillation was delayed in the presence of lipid nanodiscs lacking cholesterol (Figure 2). In the presence of both DOPC and DOPC:PE:PG nanodiscs there was almost a three-fold increase in α-Syn fibrillation lag time (Figure 2B). This observation is similar to other reports where α-Syn fibrillation is delayed by the presence of lipids ^37,38,48^. Moreover, no significant changes in the fibrillation rate were observed (Figure 2C). This suggests that the lipids effectively decrease the population of free α-Syn monomers, leaving less protein available for oligomerization and fibrillation. This is, surprisingly, also observed in the presence of zwitterionic DOPC to which α-Syn has a very low affinity. It seems that even this low affinity is sufficient to almost double lag times for the onset of fibrillation. The potential for using zwitterionic small unilamellar vesicles to inhibit the onset of α-Syn fibrillation and protect dopaminergic cells has recently been reported by Aliakbari *et al.* ^62^. It seems that α-Syn engages in a dynamic exchange with PC and PE membrane components that does not give rise to high affinity binding but is nonetheless very important for preventing nucleation with subsequent fibrillation.

When cholesterol was present in the lipid nanodiscs, there was a significant promotion of α-Syn fibrillation (Figure 2). The lag times decreased almost to zero (Figure 2B) and fibrillation rate increased to more than 13 times higher than for α-Syn alone (Figure 2C). One possible explanation for these dramatic changes is that lipid nanodiscs containing cholesterol served as a seeding agent, independently of other lipids present. Other reports on the effect of cholesterol on α-Syn fibrillation are conflicting. Some studies using lipid vesicles did not observe any promotion of α-Syn fibrillation by lipid vesicles containing cholesterol ^25,62^, while other studies observed increased oligomerization in the presence of oxidized cholesterol metabolites ^66^ or increased fibrillation and deposit formation in cell cultures exposed to higher cholesterol concentrations ^67^. The lipid nanodiscs used in our experiments are much smaller (see Table S1) than vesicles used in previous experiments and there is a lack of stored curvature elastic stress. Thus, it is possible that cholesterol can promote co-localization and co-orientation of several α-Syn molecules in a spatially restricted area. This effect of local enrichment of α-Syn on membranes and molecular crowding has been suggested as the mechanism of α-Syn fibrillation in live cells ^68^. However, a more detailed insight of how the presence of different membrane components affects the different parts of the protein is required.

Using the approach of Shortridge *et al.* ^31^ it was possible to determine aK_D_ values for individual residues of α-Syn primary, and in this way gain some insight into how different membrane components affects different parts of the protein. The aK_D_ values do not differentiate between the possible causes of line-broadening but are all relatable to the effect the presence of lipids has on the protein. While the approached yielded valuable information, we can observe and fit only 48% of the amino-acid sequence. This number is reduced for samples containing 34% cholesterol. The undetectable amino-acid states suggest that protein-lipid, protein-protein or intramolecular exchange takes place on a time-scale unfavourable for NMR observation. Notably no lysines, which have a strong electrostatic affinity for lipid head groups ^38,64^, were observed. In fact, if we inspect which amino acids gives rise to undetectable peaks and where they appear in the sequence (Figure 3B), we find that most invisible states are organized in non-perfect hexameric KTKEGV repeats which represent a small α-helical motif with high affinity to lipid environment ^38,58,69,70^. The fact that the presence of cholesterol cause more α-Syn residues in these repeats to become undetectable suggests that these are involved in cholesterol interaction. This is particularly apparent in the N-terminal region and NAC region, where we observed a longer stretch of invisible peaks in the T72-V82 region. This stretch coincides with the ending of the NAC core which has been proposed to function as a sensor of the state of the lipid bilayer ^52^, and which participates in the formation of β-sheets during oligomerization ^71^. Notably, this invisible stretch is extended when cholesterol is present, particularly in the presence of DOPC:PE:PG with 30% cholesterol nanodiscs, suggestive of a link to the seeding-like activity of nanodiscs containing cholesterol reported here. We also observed the induction of several invisible peaks in the C-terminal (especially in the interval G132-D135) which is, generally, regarded as non-reactive towards lipids but protective against aggregation ^72,73^. This could mean that these residues are involved in a chaperone-like activity, as has been suggested ^48,73^, maintaining at least some protein interaction with the solvent rather than allowing it to nucleate at the membrane or other potential seeding sites. In this scenario, the influence of cholesterol on this part of the protein would interfere with the moderating effect the C-terminal of α-Syn, possibly speeding up nucleation. Interestingly, it has recently been shown that the C-terminal helps to modulate α-Syn membrane interaction and its localization at the pre-synaptic terminal by binding to Ca^2+60^. Moreover, natively occurring C-terminally truncated α-Syn is associated with increased fibrillation, a phenomenon that was further enhanced if known disease mutants were present in the fibrillation assays ^73^.

The known mutants of α-Syn involved in hereditary early onset of PD are clustered towards the end of the membrane binding N-terminal segment of the protein ^4,51^. Of these sequence positions, only A30 and A53 could be directly observed in our results. The A53 aK_D_ was high for all lipid samples when compared, for example, to the medium affinity of V15 and low affinity of E131, suggesting that A53 have a general role in lipid binding (Figure 4C). A30, in contrast, has a low affinity, with no aK_D_ determined for the DOPC:PE:PG model with cholesterol. Its mutant A30P has been reported to decrease α-Syn binding to membranes ^64^, while A53T are associated with slower membrane-perturbing effects ^7^.

In our study we have shown that cholesterol inhibits binding of α-Syn to lipid membranes. However, we have also observed that lipid nanodiscs containing cholesterol can act as a strong promotor of α-Syn fibrillation. Both inhibition of α-Syn binding to vesicles and promotion of α-Syn fibrillation by lipid nanodiscs are seemingly independent of lipid composition as they are similar for both DOPC and DOPC:PE:PG (4:3:1) lipid models. Still, when investigated by NMR we have shown that the effect of cholesterol on α-Syn is fundamentally different for different lipid bilayers. In zwitterionic lipid models (DOPC) it is the NAC region which is affected the most by the presence of cholesterol, while in the DOPC:PE:PG lipid model it is the C-terminal and several N-terminal residues. The recently published model of Viennet *et al.* for α-Syn-lipid interaction and fibrillation were able to rationalize fibrillation in terms of lipid charge and fluidity ^4^; in this model, a paucity of high-affinity charged sites may lead to molecular crowding where NAC segments are aligned resulting in accelerated oligomerization. Relating our results to this model, we propose that cholesterol-rich regions could act as nucleation sites. If sites with net negative charge are not available, cholesterol-rich sites will become the next preferred site for binding and potential crowding. Moreover, cholesterol would make it harder for a polypeptide to intercalate into the bilayer ^9,74,75^, leaving more scope for protein-protein interaction at the membrane. This difference in α-Syn binding behaviour could be used to explain some discrepancy between observed α-Syn interactions with cholesterol, primarily by making a distinction between a cholesterol-rich, NAC-supported binding mode and a high-charge, high fluidity mode supported primarily by the N-terminal. Both modes can lead to increased fibrillation if molecular crowding occurs at the available sites, but cholesterol strongly promotes these events independently of charge. Given the high amount of cholesterol in the synaptic membrane, this suggests that α-Syn must neither overload potential nucleation sites nor remain at such sites for an extended time. A possible interaction partner of α-Syn that might alleviate such overload, are the ubiquitous 14-3-3 proteins that normally have roles as scaffolding proteins that bring other proteins together. Several of the isoforms (seven in humans) of this protein class have been identified in Lewy bodies ^76,77^. Interaction studies of the η-isoform and α-Syn showed that it binds to oligomeric, but not monomeric α-Syn, and intriguingly this isoform also are notable for its presence in the synaptic membrane and synaptic junctions ^78^. In addition, the isoform was shown to reduce α-Syn toxicity in cell models ^79^. These observations suggest that 14-3-3 proteins may be involved in handling α-Syn overload situations at cholesterol-rich sites, and a may provide a link between the lipid membrane and the proteostasis network.

## Materials and methods

### Recombinant α-Syn expression and purification

For use in this study, we modified the vector pET21a-alpha-synuclein (Addgene plasmid 51486) by introducing a 6x-histidine tag followed by a TEV cleavage site to the N-terminal of α-Syn. This was achieved with the Q5® Site-Directed Mutagenesis Kit (New England Biolabs, UK) using the primers listed in the Supplementary information (Table S2).

Unlabelled α-Syn was expressed and purified using a variant of a previously described protocol ^80^. The plasmid encoding α-Syn was transformed into E. coli BL21 Star™ (DE3) (Invitrogen, Waltham, UK) following manufacturer’s instructions and plated onto Lysogeny broth (LB) agar plates supplemented with 100 μg/mL ampicillin. Single colonies were used to inoculate 10 mL overnight starter cultures. The following day the overnight culture was used to inoculate 1 litre cultures. All the cultures were grown in media supplemented with 100μg/mL ampicillin using orbital shakers at 37 °C and 250 rpm. Protein expression was induced when the cultures reached an optical density of 0.4 (OD_620_) by the addition of IPTG to a final concentration of 0.1 mM. Cells were harvested after 5 hours by centrifugation (10 000 × g for 20 min). The obtained bacterial pellets were resuspended in 20 mL osmotic shock buffer (30 mM Tris-HCl [pH 8.0], 40% saccharose, 2 mM EDTA) per gram cell wet weight before 10 minutes incubation at room temperature. Following the osmotic shock, the cells were collected by centrifugation (20,000 × g for 20 minutes) and the content of the periplasmic space was released by adding 45 mL ice cold water containing 1 mM MgCl_2_. Cell debris was removed with centrifugation (20,000 × g for 20 minutes). The supernatant containing α-Syn was mixed with Na_2_HPO_4_ to a final concentration of 20 mM Na_2_HPO_4_ [pH 7.4] and supplemented with cOmplete™ EDTA-free protease inhibitors (Roche, Switzerland).

α-Syn was further purified with Ni-NTA affinity chromatography using a HisTrap HP 5 mL column (GE healthcare life sciences, USA) connected to an ÄKTA system (GE healthcare life sciences, USA). The Ni-NTA purification was performed using buffer A (20 mM Tris, 150 mM NaCl, 20 mM imidazole [pH 8.0]) and buffer B (20 mM Tris, 150 mM NaCl, 500 mM Imidazole [pH 8]). After elution the buffer was changed into TEV cleavage buffer (20 mM Tris, 0.5 mM EDTA, 1 mM DTT, 150 mM NaCl [pH 8.0]) using PD-10 columns (GE healthcare life sciences).

The concentration of uncleaved α-Syn was determined using an extinction coefficient at 280 nm of 7450 M^−1^ cm^−1^. TEV (produced in the house according to van den Berg *et al.* ^81^) was added to α-Syn in a 1:100 molar ratio (TEV:α-Syn) and incubated for 16 hours at 4 °C. Lastly, α-Syn was purified by size exclusion chromatography using a HiLoad Superdex column 16/600 75 pg (GE healthcare life sciences) connected to an ÄKTA system (GE healthcare life sciences). The concentration of purified α-Syn was determined using an extinction coefficient at 280 nm of 5960 M^−1^cm^−1^. Typical yield was 15 mg of cleaved protein per litre of medium.

For isotopic labelling we adapted a previously published technique in which the culture is initially grown using LB ^82,83^. The culture was grown identically to the unlabelled up to the point where the optical density reached 0.7. At this point, the cells were harvested and resuspended into M9 salt wash medium (25 mM KH_2_PO_4_, 10 mM NaCl, 5 mM MgSO_4_ 0.2 mM CaCl_2_ [pH 8.0]). Following a subsequent centrifugation (10 000 × g for 20 min) step the cells were resuspended in Modified Marley Minimal Medium (25 mM KH_2_PO_4_ [pH 8.0], 10 mM NaCl, 5mM MgSO_4_, 0.2 mM CaCl_2_, 1x Trace metal solution according Studier ^83^, 0.25x Vitamins (100x BME vitamins stock solution, SIGMA), 0.1% ^15^NH_4_Cl and 1.0% glucose or ^13^C-glucose). The medium was added in a 1:4 (v/v) ratio, i.e., for each 1 L of initial LB culture was added 250 mL of 4M media. The cultures were further grown for 1 hour at 37°C. Expression was induced by adding IPTG to a final concentration of 0.8 mM and the cells were grown for 5 hours in orbital shakers at 37 °C and 250 rpm. Further purification was identical to the unlabelled protein purification.

### Preparation of Large unilamellar vesicles

LUVs were prepared using standard methods ^84^. In brief lipid/cholesterol mixtures were prepared by dissolving defined amounts of dried lipid powders (DOPC, DOPE, DOPG and cholesterol from Avanti, USA) in 3:1 dichloromethane:methanol mixtures followed by drying using nitrogen gas, and lastly in vacuo. The lipid films were hydrated to 20 mM concentration of lipids by adding 50 mM Tris-HCl [pH 8.0]. The hydrated lipids where subsequently subjected to 4 rapid freeze-thaw cycles (−195 °C to +60 °C), and extruded (13 passes) through a 100 nm polycarbonate filter (Mini-Extruder; Avanti, USA). The sizes of vesicles were measured by dynamic laser light-scattering system (DLS) using Nanosizer ZS (Malvern Instruments, UK). The mean size of all vesicles were 100 ± 6 nm with a polydispersity index <0.1.

### Nanodisc preparation

SMA was prepared as described previously ^85^. Briefly, Xiran SZ30010 (Polyscope, Geleen, NL) was refluxed for 4 hours in 5 % (w/v) in 1 M KOH. Once cool, 6 M HCl was added dropwise to the solution at room temperature to a final concentration of 1.1 M, yielding a precipitate. The precipitate was washed 3 times with 50 mL of 100 mM HCl, then with 50 mL water. The precipitate was then freeze-dried. The loss of the anhydride groups was demonstrated by FTIR spectroscopy through the appearance of a maleic acid carbonyl stretch at 1570 cm^−1^ and the loss of the maleic anhydride carboxyl stretch at 1780 cm^−1^. The polymer was stored as a solution of 8% (w/v) SMA in 50 mM Tris-HCl (pH 8.0) at −80 °C and was used without further purification.

Nanodiscs were prepared using a modified procedure based on Scheidelaar *et al.* ^86^ Hydrated lipids and SMA were mixed in varying ratio based on nanodiscs composition (see Table S1). The mixture was processed in the same manner as hydrated lipids in LUVs preparation (4 cycles freeze-thawing and 13 extrusion cycles through a 100 nm filter), and incubated overnight at room temperature. Nanodiscs were further purified by size exclusion chromatography using HiLoad Superdex column 16/600 200 pg (GE healthcare life sciences) connected to an ÄEKTA system. The fractions containing the desired nanodiscs were collected and their size was determined by DLS using Nanosizer ZS (Malvern Instruments). The mean values of major nanodiscs peaks are shown in Table 1. The lipid concentrations of the nanodiscs was determined by taking 100 μL of the purified nanodiscs, freeze-drying them overnight (Alpha 1-2 LDplus, Christ, Germany), dissolving them in 300 μL of CUBO solvent (800 μM guanidine chloride in mixture of dimethylformamide (DMF) and trimethylamine (3:1); 20 % of DMF was d-7 deuterated; Sigma-Aldrich) containing 50 μM trimethyl phosphate standard (Sigma-Aldrich) and collect ^31^P NMR with AVANCE^TM^ III HD 600 MHz NMR instrument (Bruker, Germany).

### SPR assays

The SPR experiments were conducted using a T200 BIAcore instrument (GE Healthcare) on an L1 chip. The surface of the L1 chip was conditioned with three consecutive 1 minute injections of 20 mM CHAPS followed by one injection of 30% ethanol. The flowrate for these injections was 30 μL/min. Liposomes (10 mM in PBS) were deposit in all cells for 40 min at a flowrate of 2 μL/min. The surface was stabilized by three injections containing 100 mM NaOH for 1 min using a flowrate of 30 μL/min. The successful surface coverage was tested by injecting 0.1 mg/mL of bovine serum albumin (BSA) for 1 min at 30 μL/min, and a change of less than 400 RU indicated sufficient coverage. Between experiments, the chip surface was cleaned by subsequent injection of 20 mM CHAPS, 40 mM Octyl β-D-glucopyranoside, and 30% ethanol, each for 1 min at 30 μL/min.

For all measurement, the flow rate was fixed at 15 μL/min. Dilutions of α-Syn (from 8 μM to 0.063 μM) were injected over immobilized liposomes on the L1 chip. Injections were made from low to high concentration with 250 sec contact time and 400 sec dissociation phase. Liposomes were then regenerated by three subsequent injections of 100 mM NaOH at 30 μL/min for 1 min. The RU stability was checked and experiments were only executed if the RU was in the range of ±200 prior to the α-Syn injection. The control flow cell was treated the same way as sample cells (liposomes coverage, stabilization, BSA coverage, regeneration and cleaning) except that α-Syn injections were replaced with buffer (20 mM sodium phosphate [pH 6], 100 mM NaCl). Control flow cell background was subtracted from the experimental cell before further data processing. The data points for steady state affinity analysis were taken from 240 sec after injections. The resulting curves were fitted into Equation 1 using in-lab written Matlab script (Matlab R2017b).

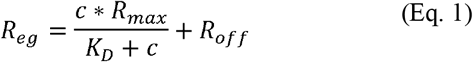

**Equation 1**: Steady-state SPR interaction fitting model where R_eg_ = response at equilibrium, c = concentration of α-Syn, R_max_ = maximum response, K_D_ = dissociation constant and R_off_ = response offset.

### ThT and TPE-TPP assay

The aggregation experiments of α-Syn were performed using a FLUOstar Optima microplate fluorescence reader (BMG Labtech, Germany) and monitored by Thioflavin T (ThT, Sigma-Aldrich) and TPE-TPP (kind gift from Professor Ben Zhong Tang ^32^) fluorescence. The fluorescence was measured at time intervals of 600 sec with 300 sec, 3 mm orbital shaking before measuring for 150 hours. The excitation and emission wavelengths were 430(10) nm and 485(5) nm for ThT and 360(10) nm and 420(10) nm for TPE-TPP. Lamp settings were: 10 flashes per well and 1300 gain for ThT and 1900 gain for TPE-TPP. Experiments were performed using black 384-well plate with clear bottom (Corning, USA, Cat. N. 3762) sealed with sealing tape (ThermoFisher Scientific, USA, Cat. N. 232701) to prevent evaporation. The aggregation buffer contained 20 mM sodium phosphate [pH 6], 100 mM NaCl, 10 mM NaN_3_, and 1 mM EDTA. The final concentrations of the components in the wells were 60 μM α-Syn, 5 μM ThT or TPE-TPP, and none or 1 mM of the lipid nanodiscs. Buffer and lipids alone were also measured as controls. We performed three independent experiments with 8-12 repeats for each sample. The recorded curves were processed using in-lab written Matlab script (Matlab R2017b). Similar to the work of Aftiska *et al.* ^48^, cumulative curves were recorded evaluating the percentage of wells having a signal three times higher than noise (mean of control wells containing all the components except α-Syn). Each curve was also fitted with Finke-Watzky two-step model Equation 2,3 and t_n_ and ν was calculated ^49^.

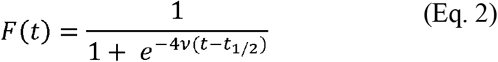

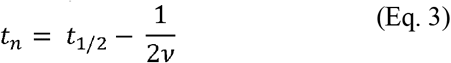

**Equation 2 and 3**: The Finke-Watzky two-step model ^49^, where t_1/2_ is a half time of fibrillation, ν is growth rate and t_N_ is lag time.

### Atomic Force Microscopy

Samples for AFM analysis were prepared in the following way. 2 uL of sample solution was deposited on a freshly cleaved mica disc and left for 30 min. in a closed Petri dish (plastic, 50 mm in diameter). After deposition, 100 uL of Milli-Q-water (filtered and de-ionized 18.2 MΩ·cm at 25 °C) was added to each mica disc, incubated for one min with subsequent excess water removal with paper tissues. This procedure was applied three times and samples were left to dry in a closed Petri dish for two hours. Sample morphologies were studied using a Bruker Bioscope Catalyst Microscope in Peak force Quantitative Nanomechanical Property Mapping mode. Silicon Nitride Bruker Scanasyst Air cantilevers with ~ 2 nm tip radius and ~ 25 ° tip angle were used for imaging. Images were taken at 256px · 256 px resolution. At least five images were taken for each sample, which were at least 50 um apart from each other to ensure that the morphologies shown are representative.

### NMR spectroscopy

NMR spectra of 0.1 μM uniformly labelled ^15^N α-Syn in 20 mM sodium phosphate [pH 6], 100 mM NaCl and 10% D_2_O were measured at 299K using 850 MHz (Bruker, Germany). We used best-HSQC(b_hsqcetf3gpsi)^94^ and DOSY (stebpgls19) standard pulse programs supplied in TopSpin 4.0.1. Spectra were recorded in the absence or presence of lipid nanodiscs at 500, 250, 125, or 62.5 μM total lipid concentration. Data were processed using TopSpin, PINT ^87^ and in-lab written Matlab scripts (Matlab 2017b). Apparent K_d_ (aK_d_) calculations were done according to Shortridge *et al.* ^31^, the area of deconvoluted peaks from PINT was used to calculate peak volume ratio B = 1 − (bound peak volume/free peak volume) and fitted according Equation 4.

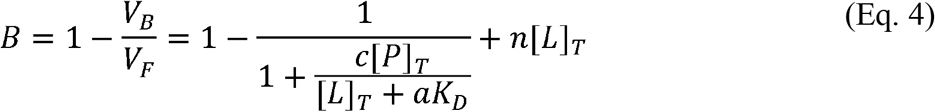

**Equation 4**: Protein-ligand binding model based on Shortridge *et al.* ^31^ where B is the NMR volume ratio, V_B_ the peak volume in bounded state, VF the peak volume in free state, c the unitless NMR area ratio constant, [P]_T_ the total protein concentration, [L]_T_ is the total lipid concentration, aK_D_ is the apparent dissociation constant, and n is the nonspecific binding constant.

## Acknowledgements

This research was funded by The Norwegian Research Council grant NFR240063 and partially supported through the Norwegian NMR Platform, NNP (infrastructure grant 226244/F50). The authors would like to thank the staff at the NNP Bergen node for facilitating the NMR experimental work, Zhang *et al.* for providing TPE-TPP, and Stefan Scheidelaar at Polyscope and Jonas Dörr at University of Utrecht for providing SMA.

## Author contributions

ØH and MJ conceived the research questions. ØH, MJ and SF designed the experiments. Lipid nanodiscs were established by MJ and SF. α-Syn expression was established by ML. SPR data was collected and analysed by MJ and DT. NMR data were collected and analysed by MJ, EB, ØH. All other laboratory work was carried out by MJ, EB, II and VG. All authors interpreted data and commented on the final version of the manuscript. The original grant proposal and the research program was conceived, designed and written by ØH.

## References

1. Winner, B. et al. In vivo demonstration that alpha-synuclein oligomers are toxic. Proc. Natl. Acad. Sci. U. S. A. 108, 4194–4199 (2011).

2. Logan, T., Bendor, J., Toupin, C., Thorn, K. & Edwards, R. H. α-Synuclein promotes dilation of the exocytotic fusion pore. Nat. Neurosci. 20, 681–689 (2017).

3. Galvagnion, C. The Role of Lipids Interacting with α-Synuclein in the Pathogenesis of Parkinson’s Disease. Journal of Parkinson’s Disease (2017). doi:10.3233/JPD-171103

4. Viennet, T. et al. Structural insights from lipid-bilayer nanodiscs link α-Synuclein membrane-binding modes to amyloid fibril formation. Commun. Biol. 1, 44 (2018).

5. Jo, E., McLaurin, J. A., Yip, C. M., St. George-Hyslop, P. & Fraser, P. E. α-Synuclein membrane interactions and lipid specificity. J. Biol. Chem. 275, 34328–34334 (2000).

6. Galvagnion, C. et al. Lipid vesicles trigger α-synuclein aggregation by stimulating primary nucleation. Nat. Chem. Biol. 11, 229–234 (2015).

7. Middleton, E. R. & Rhoades, E. Effects of curvature and composition on α-synuclein binding to lipid vesicles. Biophys. J. (2010). doi:10.1016/j.bpj.2010.07.056

8. van Meer, G. & de Kroon, A. I. P. M. Lipid map of the mammalian cell. J. Cell Sci. 124, 5–8 (2011).

9. Róg, T., Pasenkiewicz-Gierula, M., Vattulainen, I. & Karttunen, M. Ordering effects of cholesterol and its analogues. Biochim. Biophys. Acta – Biomembr. 1788, 97–121 (2009).

10. Robinson, A. J., Richards, W. G., Thomas, P. J. & Hann, M. M. Behavior of cholesterol and its effect on head group and chain conformations in lipid bilayers: a molecular dynamics study. Biophys. J. 68, 164–170 (1995).

11. Kojro, E., Gimpl, G., Lammich, S., Marz, W. & Fahrenholz, F. Low cholesterol stimulates the nonamyloidogenic pathway by its effect on the alpha-secretase ADAM 10. Proc. Natl. Acad. Sci. 98, 5815–5820 (2001).

12. Simons, M. et al. Cholesterol depletion inhibits the generation of beta-amyloid in hippocampal neurons. Proc. Natl. Acad. Sci. U. S. A. 95, 6460–4 (1998).

13. Valenza, M. et al. Dysfunction of the Cholesterol Biosynthetic Pathway in Huntington’s Disease. J. Neurosci. 25, 9932–9939 (2005).

14. Vance, J. E. Transfer of cholesterol by the NPC team. Cell Metab. 12, 105–106 (2010).

15. Wang, M. L. et al. Identification of surface residues on Niemann-Pick C2 essential for hydrophobic handoff of cholesterol to NPC1 in lysosomes. Cell Metab. 12, 166–173 (2010).

16. Wood, W. G., Igbavboa, U., Müller, W. E. & Eckert, G. P. Cholesterol asymmetry in synaptic plasma membranes. Journal of Neurochemistry (2011). doi:10.1111/j.1471-4159.2010.07017.x

17. Arenas, F., Garcia-Ruiz, C. & Fernandez-Checa, J. C. Intracellular Cholesterol Trafficking and Impact in Neurodegeneration. Front. Mol. Neurosci. 10, (2017).

18. Egawa, J., Pearn, M. L., Lemkuil, B. P., Patel, P. M. & Head, B. P. Membrane lipid rafts and neurobiology: age-related changes in membrane lipids and loss of neuronal function. J. Physiol. 594, 4565–4579 (2016).

19. Hu, G., Antikainen, R., Jousilahti, P., Kivipelto, M. & Tuomilehto, J. Total cholesterol and the risk of Parkinson disease. Neurology 70, 1972–1979 (2008).

20. Powers, K. M. et al. Dietary fats, cholesterol and iron as risk factors for Parkinson’s disease. Parkinsonism Relat. Disord. 15, 47–52 (2009).

21. Huang, X. et al. Serum cholesterol and the progression of parkinson’s disease: Results from DATATOP. PLoS One 6, e22854 (2011).

22. Di Scala, C. et al. Common molecular mechanism of amyloid pore formation by Alzheimer’s β-amyloid peptide and α-synuclein. Sci. Rep. (2016). doi:10.1038/srep28781

23. Fantini, J. et al. Bexarotene Blocks Calcium-Permeable Ion Channels Formed by Neurotoxic Alzheimer’s beta-Amyloid Peptides. Acs Chem. Neurosci. 5, 216–224 (2014).

24. Fantini, J., Carlus, D. & Yahi, N. The fusogenic tilted peptide (67–78) of α-synuclein is a cholesterol binding domain. Biochim. Biophys. Acta – Biomembr. 1808, 2343–2351 (2011).

25. O’Leary, E. I. et al. Effects of phosphatidylcholine membrane fluidity on the conformation and aggregation of N-terminally acetylated α-synuclein. J. Biol. Chem. jbc.RA118.002780 (2018). doi:10.1074/jbc.RA118.002780

26. Shvadchak, V. V., Falomir-Lockhart, L. J., Yushchenko, D. A. & Jovin, T. M. Specificity and Kinetics of alpha-Synuclein Binding to Model Membranes Determined with Fluorescent Excited State Intramolecular Proton Transfer (ESIPT) Probe. J. Biol. Chem. 286, 13023–13032 (2011).

27. Sezgin, E. & Schwille, P. Model membrane platforms to study protein-membrane interactions. Mol. Membr. Biol. (2012). doi:10.3109/09687688.2012.700490

28. Borch, J. & Hamann, T. The nanodisc: A novel tool for membrane protein studies. Biol. Chem. 390, 805–814 (2009).

29. Dörr, J. M. et al. Detergent-free isolation, characterization, and functional reconstitution of a tetrameric K ^+^ channel: The power of native nanodiscs. Proc. Natl. Acad. Sci. 111, 18607–18612 (2014).

30. Denisov, I. G., Grinkova, Y. V., Lazarides, A. A. & Sligar, S. G. Directed Self-Assembly of Monodisperse Phospholipid Bilayer Nanodiscs with Controlled Size. J. Am. Chem. Soc. (2004). doi:10.1021/ja0393574

31. Shortridge, M. D., Hage, D. S., Harbison, G. S. & Powers, R. Estimating Protein-Ligand Binding Affinity using High- Throughput Screening by NMR. J Comb Chem 10, 948–958 (2009).

32. Leung, C. W. T. et al. Detection of oligomers and fibrils of α-synuclein by AIEgen with strong fluorescence. Chem. Commun. 51, 1866–1869 (2015).

33. Vorobyov, I. & Allen, T. W. On the role of anionic lipids in charged protein interactions with membranes. Biochim. Biophys. Acta – Biomembr. (2011). doi:10.1016/j.bbamem.2010.11.009

34. Yeung, T. et al. Membrane phosphatidylserine regulates surface charge and protein localization. Science (80−.). 319, 210–213 (2008).

35. Stahelin, R. V., Scott, J. L. & Frick, C. T. Cellular and molecular interactions of phosphoinositides and peripheral proteins. Chem. Phys. Lipids (2014). doi:10.1016/j.chemphyslip.2014.02.002

36. Rødland, I., Halskau, Ø., Martínez, A. & Holmsen, H. α-Lactalbumin binding and membrane integrity – Effect of charge and degree of unsaturation of glycerophospholipids. Biochim. Biophys. Acta – Biomembr. 1717, 11–20 (2005).

37. Grey, M., Linse, S., Nilsson, H., Brundin, P. & Sparr, E. Membrane interaction of α-synuclein in different aggregation states. J. Parkinsons. Dis. 1, 359–371 (2011).

38. Pirc, K. & Ulrih, N. P. α-Synuclein interactions with phospholipid model membranes: Key roles for electrostatic interactions and lipid-bilayer structure. Biochim. Biophys. Acta – Biomembr. 1848, 2002–2012 (2015).

39. Nuscher, B. et al. α-synuclein has a high affinity for packing defects in a bilayer membrane: A thermodynamics study. J. Biol. Chem. 279, 21966–21975 (2004).

40. Rhoades, E., Ramlall, T. F., Webb, W. W. & Eliezer, D. Quantification of α-synuclein binding to lipid vesicles using fluorescence correlation spectroscopy. Biophys. J. 90, 4692–4700 (2006).

41. Zuidam, N. J., Gouw, H. K. M. E., Barenholz, Y. & Crommelin, D. J. A. Physical (in) stability of liposomes upon chemical hydrolysis: the role of lysophospholipids and fatty acids. BBA – Biomembr. (1995). doi:10.1016/0005-2736(95)00180-5

42. Galvagnion, C. et al. Chemical properties of lipids strongly affect the kinetics of the membrane-induced aggregation of α-synuclein. Proc. Natl. Acad. Sci. 113, 7065–7070 (2016).

43. Van Maarschalkerweerd, A., Vetri, V. & Vestergaard, B. Cholesterol facilitates interactions between α-synuclein oligomers and charge-neutral membranes. FEBS Lett. 589, 2661–2667 (2015).

44. Biancalana, M. & Koide, S. Molecular mechanism of Thioflavin-T binding to amyloid fibrils. Biochim. Biophys. Acta – Proteins Proteomics 1804, 1405–1412 (2010).

45. Amdursky, N., Erez, Y. & Huppert, D. Molecular rotors: What lies behind the high sensitivity of the thioflavin-T fluorescent marker. Acc. Chem. Res. 45, 1548–1557 (2012).

46. Meisl, G. et al. Molecular mechanisms of protein aggregation from global fitting of kinetic models. Nat. Protoc. 11, 252–272 (2016).

47. Voropai, E. S. et al. Spectral properties of thioflavin T and its complexes with amyloid fibrils. J. Appl. Spectrosc. 70, 868–874 (2003).

48. Afitska, K., Fucikova, A., Shvadchak, V. V. & Yushchenko, D. A. Modification of C Terminus Provides New Insights into the Mechanism of α-Synuclein Aggregation. Biophys. J. 113, 2182–2191 (2017).

49. Morris, A. M., Watzky, M. A., Agar, J. N. & Finke, R. G. Fitting neurological protein aggregation kinetic data via a 2-step, minimal/“Ockham’s razor” model: The Finke-Watzky mechanism of nucleation followed by autocatalytic surface growth. Biochemistry 47, 2413–2427 (2008).

50. Bermel, W. et al. Protonless NMR experiments for sequence-specific assignment of backbone nuclei in unfolded proteins. J. Am. Chem. Soc. 128, 3918–3919 (2006).

51. Porcari, R. et al. The H50Q mutation induces a 10-fold decrease in the solubility of α-synuclein. J. Biol. Chem. 290, 2395–2404 (2015).

52. Fusco, G. et al. Direct observation of the three regions in α-synuclein that determine its membrane-bound behaviour. Nat. Commun. 5, 3827 (2014).

53. Liang, B. & Tamm, L. K. Solution NMR of SNAREs, complexin and α-synuclein in association with membrane-mimetics. Prog. Nucl. Magn. Reson. Spectrosc. 105, 41–53 (2018).

54. Terakawa, M. S. et al. Membrane-induced initial structure of α-synuclein control its amyloidogenesis on model membranes. Biochim. Biophys. Acta – Biomembr. 1860, 757–766 (2018).

55. Niklasson, M. et al. Comprehensive analysis of NMR data using advanced line shape fitting. J. Biomol. NMR 69, 93–99 (2017).

56. Davidson, W. S., Jonas, A., Clayton, D. F. & George, J. M. Stabilization of α-Synuclein secondary structure upon binding to synthetic membranes. J. Biol. Chem. 273, 9443–9449 (1998).

57. Eliezer, D., Kutluay, E., Bussell, R. & Browne, G. Conformational properties of alpha-synuclein in its free and lipid-associated states. J. Mol. Biol. 307, 1061–1073 (2001).

58. Beyer, K. Mechanistic aspects of Parkinson’s disease: α-synuclein and the biomembrane. Cell Biochem. Biophys. 47, 285–299 (2007).

59. Fantini, J. & Yahi, N. Molecular insights into amyloid regulation by membrane cholesterol and sphingolipids: Common mechanisms in neurodegenerative diseases. Expert Rev. Mol. Med. 12, 1–22 (2010).

60. Lautenschläger, J. et al. C-terminal calcium binding of α-synuclein modulates synaptic vesicle interaction. Nat. Commun. 9, 712 (2018).

61. Terakawa, M. S. et al. Impact of membrane curvature on amyloid aggregation. Biochimica et Biophysica Acta – Biomembranes (2018). doi:10.1016/j.bbamem.2018.04.012

62. Aliakbari, F. et al. The potential of zwitterionic nanoliposomes against neurotoxic alpha-synuclein aggregates in Parkinson’s Disease. Nanoscale (2018). doi:10.1039/C8NR00632F

63. Gentry, K. A. et al. Kinetic and Structural Characterization of the Effects of Membrane on the Complex of Cytochrome b 5 and Cytochrome c. Sci. Rep. (2017). doi:10.1038/s41598-017-08130-7

64. Stöckl, M., Fischer, P., Wanker, E. & Herrmann, A. α-Synuclein Selectively Binds to Anionic Phospholipids Embedded in Liquid-Disordered Domains. J. Mol. Biol. 375, 1394–1404 (2008).

65. Furse, S. et al. Evidence that Listeria innocua modulates its membrane’s stored curvature elastic stress, but not fluidity, through the cell cycle. Sci. Rep. 7, 1–11 (2017).

66. Bosco, D. A. et al. Elevated levels of oxidized cholesterol metabolites in Lewy body disease brains accelerate α-synuclein fibrilization. Nat. Chem. Biol. 2, 249–253 (2006).

67. Bar-On, P. et al. Statins reduce neuronal α-synuclein aggregation in in vitro models of Parkinson’s disease. J. Neurochem. 105, 1656–1667 (2008).

68. Peng, C. et al. Cellular milieu imparts distinct pathological α-synuclein strains in α-synucleinopathies. Nature 557, 558–563 (2018).

69. George, J. M., Jin, H., Woods, W. S. & Clayton, D. F. Characterization of a novel protein regulated during the critical period for song learning in the zebra finch. Neuron 15, 361–372 (1995).

70. Bussell, R. & Eliezer, D. A structural and functional role for 11-mer repeats in α-synuclein and other exchangeable lipid binding proteins. J. Mol. Biol. 329, 763–778 (2003).

71. Rodriguez, J. A. et al. Structure of the toxic core of α-synuclein from invisible crystals. Nature 525, 486–490 (2015).

72. Kim, T. D., Paik, S. R. & Yang, C. H. Structural and functional implications of C-terminal regions of alpha-synuclein. Biochemistry 41, 13782–13790 (2002).

73. Li, W. et al. Aggregation promoting C-terminal truncation of alpha-synuclein is a normal cellular process and is enhanced by the familial Parkinson’s disease-linked mutations. Proc. Natl. Acad. Sci. 102, 2162–2167 (2005).

74. Wennberg, C. L., Van Der Spoel, D. & Hub, J. S. Large influence of cholesterol on solute partitioning into lipid membranes. J. Am. Chem. Soc. 134, 5351–5361 (2012).

75. Alwarawrah, M., Dai, J. & Huang, J. A molecular view of the cholesterol condensing effect in DOPC lipid bilayers. J. Phys. Chem. B 114, 7516–7523 (2010).

76. Berg, D., Riess, O. & Bornemann, A. Specification of 14-3-3 proteins in Lewy bodies. Ann. Neurol. 54, 135 (2003).

77. Wakabayashi, K., Umahara, T., Hirokawa, K., Hanyu, H. & Uchihara, T. 14-3-3 protein sigma isoform co-localizes with phosphorylated α-synuclein in Lewy bodies and Lewy neurites in patients with Lewy body disease. Neurosci. Lett. 674, 171–175 (2018).

78. Martin, H., Rostas, J., Patel, Y. & Aitken, A. Subcellular Localisation of 14-3-3 Isoforms in Rat Brain Using Specific Antibodies. J. Neurochem. 63, 2259–2265 (2002).

79. Plotegher, N. et al. The chaperone-like protein 14-3-3h interacts with human a-synuclein aggregation intermediates rerouting the amyloidogenic pathway and reducing a-synuclein cellular toxicity. Hum. Mol. Genet. 23, 5615–5629 (2014).

80. Caldinelli, L., Albani, D. & Pollegioni, L. One single method to produce native and Tat-fused recombinant human alpha-synuclein in Escherichia coli. Bmc Biotechnol. 13, (2013).

81. van den Berg, S., Lofdahl, P. A., Hard, T. & Berglund, H. Improved solubility of TEV protease by directed evolution. J. Biotechnol. 121, 291–298 (2006).

82. Marley, J., Lu, M. & Bracken, C. A method for efficient isotopic labeling of recombinant proteins. J. Biomol. Nmr 20, 71–75 (2001).

83. Studier, F. W. Protein production by auto-induction in high-density shaking cultures. Protein Expr. Purif. 41, 207–234 (2005).

84. Torchilin, V. & Weissig, V. Liposomes◻: A practical approach. (Oxford University Press, 2003).

85. Swainsbury, D. J. K., Scheidelaar, S., van Grondelle, R., Killian, J. A. & Jones, M. R. Bacterial Reaction Centers Purified with Styrene Maleic Acid Copolymer Retain Native Membrane Functional Properties and Display Enhanced Stability. Angew. Chemie-International Ed. 53, 11803–11807 (2014).

86. Scheidelaar, S. et al. Molecular Model for the Solubilization of Membranes into Nanodisks by Styrene Maleic Acid Copolymers. Biophys. J. 108, 279–290 (2015).

87. Ahlner, A., Carlsson, M., Jonsson, B. H. & Lundström, P. PINT: A software for integration of peak volumes and extraction of relaxation rates. J. Biomol. NMR 56, 191–202 (2013).

