## Supplementary Information for Cholesterol is a strong promoter for "Cholesterol is a strong promotor of an α-Synuclein membrane binding mode that accelerates oligomerization" (from another version of this preprint: version 2, 2019-08-08)

#### Preparation of cholesterol-containing nanodiscs using styrene maleic acid

The DOPC nanodiscs were made using 100 nm vesicles by adding SMA to 1% *w/v*, and overnight incubation followed by SEC purification. However, this method was not suitable for the more complex lipid mixtures where 1% SMA only led to an increase in the size of the vesicles and not to nanodiscs formation (vesicle size increased from 100 nm to about 200 nm, Figure S1B). Apparently, cholesterol and PE have a negative effect on nanodisc formation. It is possible that both PE and cholesterol are able to relive packing stress which leads to more stable vesicles and therefore the transfer of lipids to the discs is, overall, less thermodynamically favourable. To ensure proper nanodisc formation, we tested several SMA concentrations until we found one with a single well-defined peak during SEC purification (Figure S1C). We also adjusted the nanodiscs preparation method by adding SMA to lipid mixtures before vesicles formation, as this returned higher yields (Table S1). We observed that nanodiscs containing cholesterol were consistently larger than those without (Figure S1D), while a further increase of SMA concentration caused only a higher variability of nanodisc size without decreasing the average dimensions (Figure S1C, peak broadening of SEC traces). SMA concentrations used for the preparation of nanodisc and the sizes of the nanodiscs formed are presented in Table 1. Dynamic light scattering (DLS) indicated no changes in the size distribution even after a week of storage at 4 °C, indicating that the nanodiscs were stable. The incorporation of cholesterol into nanodiscs was verified by LC-MS/MS (Figure S3).

| Lipid composition | SMA concentration [%] | Major nanodiscs size fraction [nm] | Average mass fraction of lipids converted to nanodiscs [%] |
|---|---|---|---|
| DOPC | 1 | 10.45 ± 2.80 | 20.9 ± 0.8 |
| DOPC with 30% cholesterol | 2.5 | 31.95 ± 10.45 | 8.9 ± 2.0 |
| DOPC:PE:PG | 2.5 | 13.50 ± 3.20 | 14.6 ± 1.7 |
| DOPC:PE:PG with 30% cholesterol | 3.5 | 28.60 ± 11.10 | 13.4 ± 1.8 |

**Table S1:** Table of lipid nanodiscs used throughout the study. SMA concentration used to prepare the nanodiscs, their size and average yield differ, based on the composition of starting vesicles. Nanodisc sizes were determined by DLS, and mass fractions where determined based on MS data.

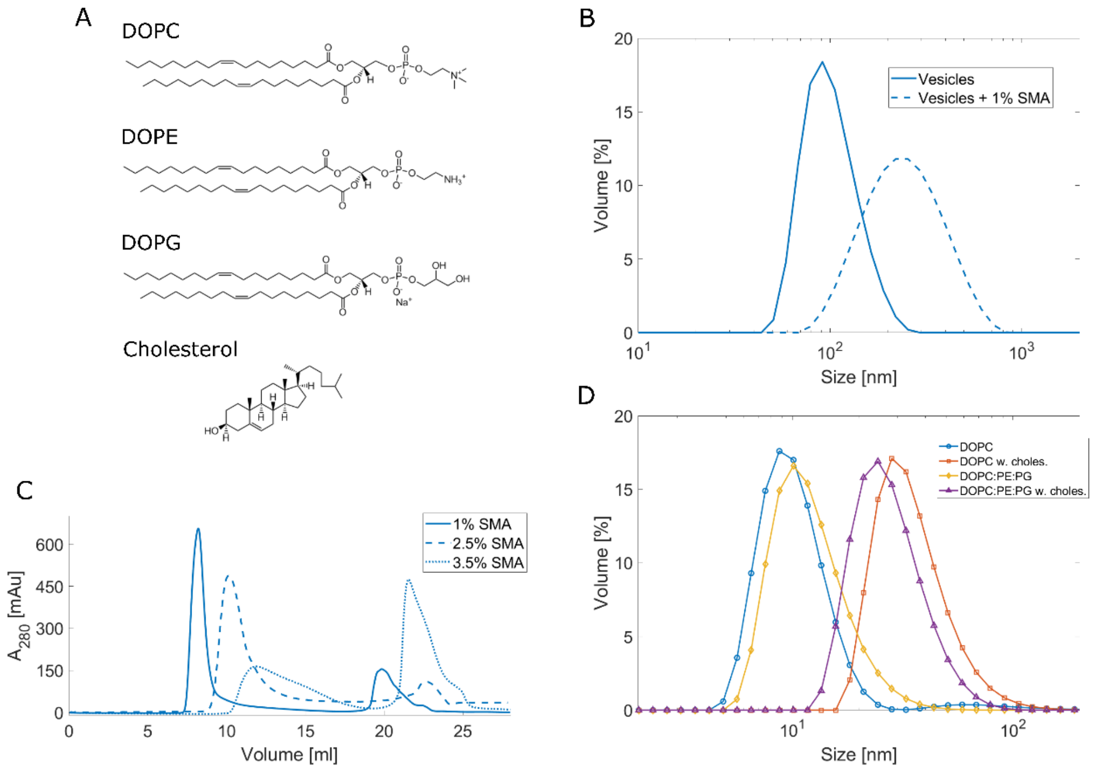

**Figure S1:** Nanodiscs preparation. (A) Structures of lipids used in nanodiscs preparation. in this work four compositions of nanodiscs have been used: DOPC, DOPC with 30% (w/w) cholesterol, DOPC:PE:PG (4:3:1) and DOPC:PE:PG with 30% (w/w) cholesterol. (B) DLS measurement of DOPC with 30% cholesterol vesicles without and with 1% SMA after overnight incubation in RT. Size of vesicles almost doubled from 100 nm to 200 nm, however disc formation was not observed. (C) SEC chromatography of DOPC with 30% cholesterol resulted in nanodisc formation after overnight incubation in RT with 1, 2.5, or 3% SMA. Leftmost peaks correspond to 200 nm vesicles at the end of dead column volume, followed by two nanodisc peaks and lastly (> 18 mL) excess of SMA not interacting with vesicles is eluted. (D) DLS measurement of prepared nanodiscs which were purified by SEC. Both DOPC and DOPC:PE:PG (4:3:1) nanodiscs without cholesterol are smaller (10 and 13 nm) than nanodiscs containing 30% (w/w) cholesterol (32 and 29 nm).

### Monomeric state of α-Synuclein

The stability of monomers and low-molecular-weight oligomers of α-Synuclein (α-Syn) depends on its environment, including purification and storage conditions. Oligomerization rates are affected by metal ions [1], salts [2] , temperature [3] and the α-Syn concentration itself [4]. To ensure that the α-Syn used for these experiments were kept in a monomeric state, samples were frozen immediately after the last purification step and stored at -80 °C. Freeze-drying before storage, common laboratory practice for handling some amyloid proteins, was not employed as we observed the formation of higher molecular weight oligomers during this treatment, in agreement with recently published work [5]. After thawing, the oligomeric state of our stock solutions was investigated using SEC, proton diffusion optimized NMR spectroscopy (DOSY), and by ThT and TPE-TPP fluorescence (Figure S2A, 2B and 2C).

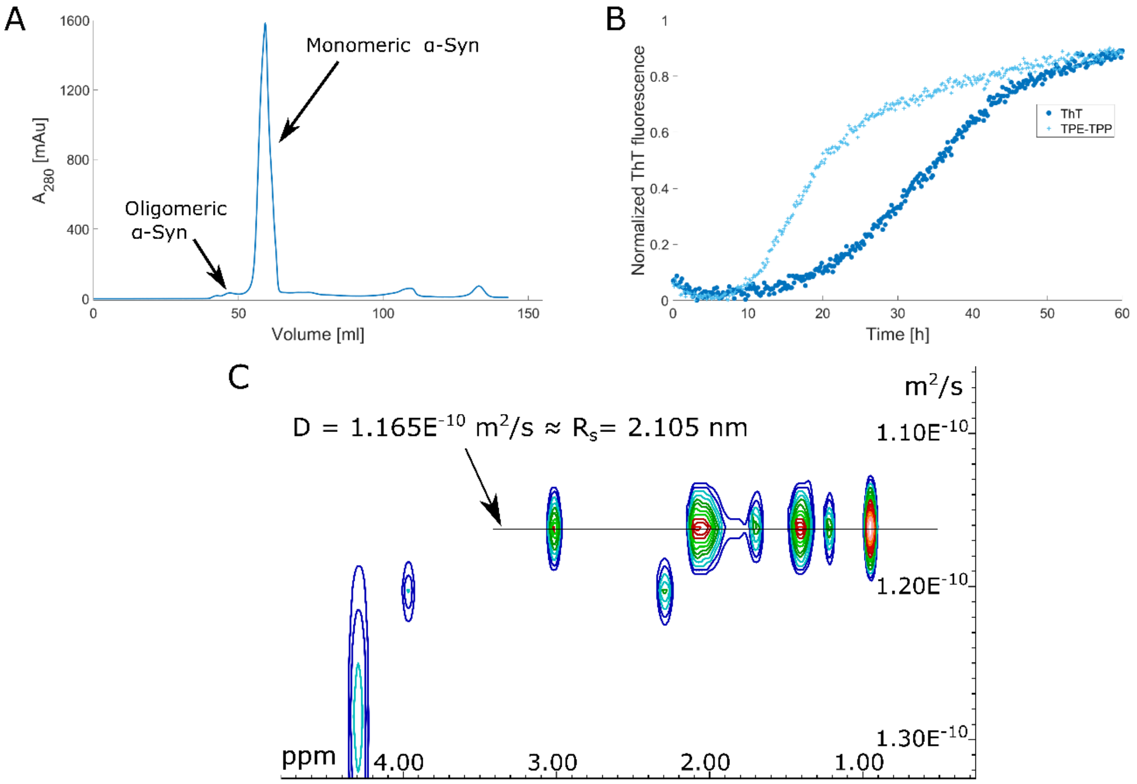

**Figure S2:** Monomeric α-Syn. (A) SEC purification of α-Syn. Major peak is monomeric α-Syn. (B) ThT experiment shows that fibrillation of α-Syn start only after 20 hours of shaking at 37 °C. TPE-TPP (more sensitive fluorescent probe) show the first sign of fibrillation after 10 hours of shaking. Both of these results suggest that the starting state of α-Syn was monomeric. (C) Example of DOSY α-Syn measurement. $^{1}$H peaks in the range from 0.8 to 3.1 ppm has a diffusion time of $1.65{\cdot}10^{-10}$ which corresponds roughly to $R_s$ (hydrodynamic radius) of 2.1 nm. This suggests that α-Syn is in a monomeric unfolded state.

[1] Paik *et al., Biochem. J.* **340**, 821 (1999)
[2] Munishkina *et al., Biochemistry* **43**, 3289–3300 (2004)
[3] Uversky *et al., J. Biol. Chem.* **276**, 10737–10744 (2001)
[4] Uversky *et al., FEBS Lett.* **515**, 99–103 (2002)
[5] Stephens *at al., Anal. Chem* **90**, 6975-6983 (2018)

### LC-MS/MS analysis of nanodiscs for the presence of cholesterol

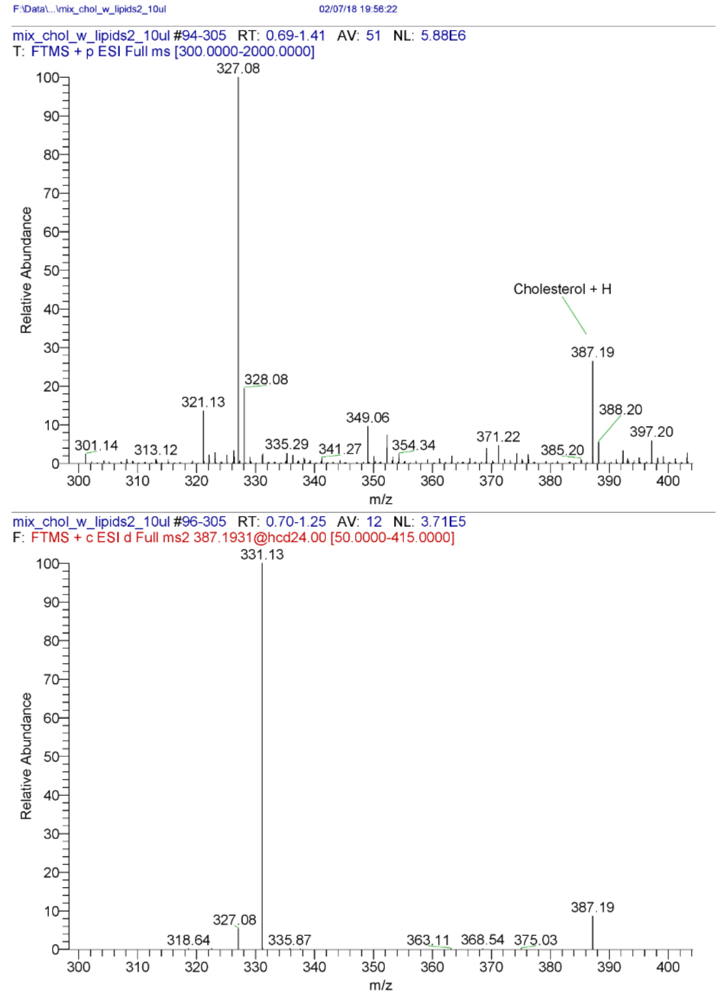

**Figure S3:** LC-MS/MS analysis of nanodiscs for the presence of cholesterol. The upper panel contains the MS1 spectrum. The lower panel contains the MS2 spectra of 387.17 m/z peak specific for cholesterol. The presence of the 331.13 m/z signal confirms identification of cholesterol.

### TPE-TPP fluorescence is inhibited by the presence of lipids.

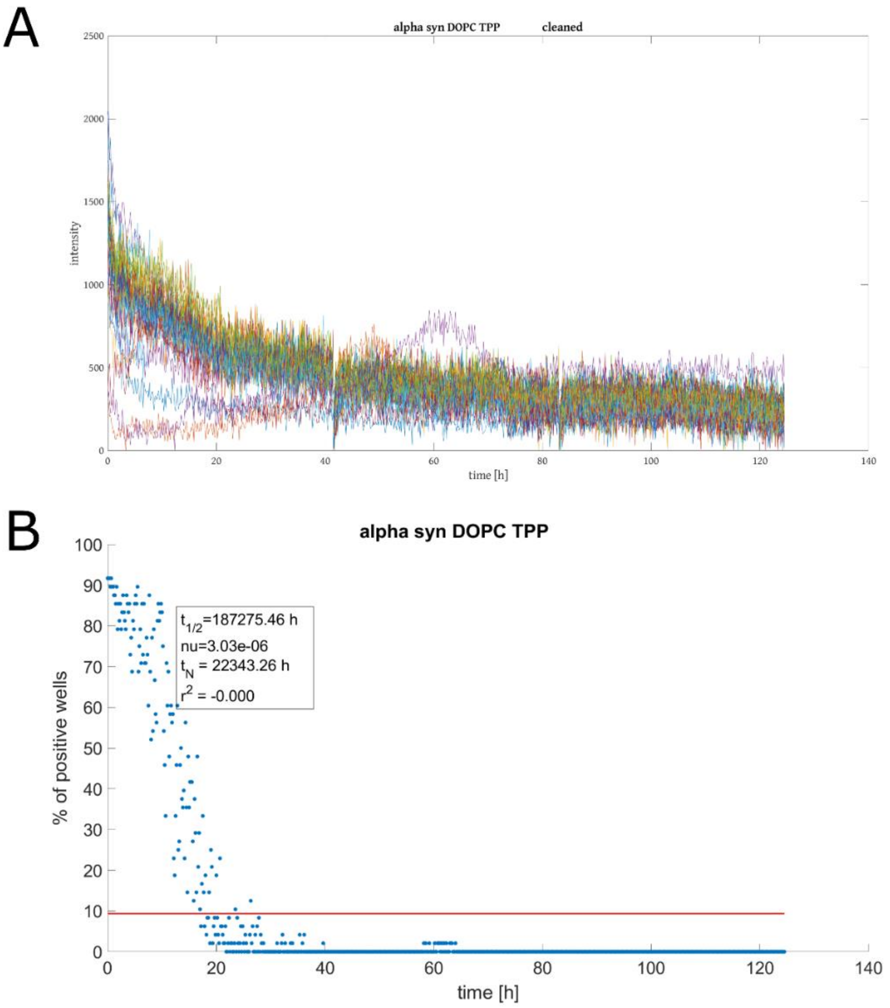

**Figure S4:** TPE-TPP fluorescence is inhibited by the presence of lipids. (A) Example of fibrillation experiment for α-Syn in the presence of DOPC nanodiscs, monitored by TPE-TPP. (B) Cumulative curve evaluation of panel (A), except for high noise on the beginning of incubation there is no increase of fluorescence later on. Similar results have been observed for all the lipid compositions used in experiments.

### Primers used for site-directed mutagenesis

| Name (F/R) | Oligo (Uppercase = target-specific primer) | Len | % GC | Tm | Ta * |
|---|---|---|---|---|---|
| a-syn_His_TEV_F | tcttctgggatcgaggaaaacctgtacttccaaATGGATGTGTTTATGAAAGG | 53 | 42 | 55°C | 58°C |
| a-syn_His_TEV_R | accatggtgatggtgatggtgagaagaacccatATGTATATCTCCTTCTTAAAGTTAAAC | 60 | 38 | 57°C | |

**Table S2:** Primers used for site-directed mutagenesis addition of His tag and TEV cleavage site on N-terminal of α-Syn in Addgene plasmid 51486.

**Residue specific $aK_D$ values plotted in heat map in Figure 3B**

| Residue specific $aK_D$ values plotted in heat map in Figure 5B. **Residue** | **DOPC** | **DOPC w. 30% choles** | **DOPC:PE:PG** | **DOPC:PE:PG w. 30% choles** | **Residue** | **DOPC** | **DOPC w. 30% choles** | **DOPC:PE:PG** | **DOPC:PE:PG w. 30% choles** | **Residue** | **DOPC** | **DOPC w. 30% choles** | **DOPC:PE:PG** | **DOPC:PE:PG w. 30% choles** |
|---|---|---|---|---|---|---|---|---|---|---|---|---|---|---|
| **1** | NaN | NaN | NaN | NaN | **26** | 45.18 | 40.67 | 45.57 | 99.14 | **51** | NaN | NaN | NaN | NaN |
| **2** | NaN | NaN | NaN | NaN | **27** | 19.03 | 9.82 | 12.44 | NaN | **52** | 90.27 | 40.49 | 81.50 | 84.82 |
| **3** | 50.14 | 20.50 | 42.94 | 38.92 | **28** | 18.86 | 10.47 | 13.40 | NaN | **53** | 35.74 | 21.68 | 27.98 | 37.13 |
| **4** | 5.29 | 4.08 | 5.41 | 1.95 | **29** | 14.66 | NaN | 10.22 | 20.10 | **54** | 45.33 | 11.75 | 31.55 | 41.47 |
| **5** | NaN | NaN | NaN | NaN | **30** | Inf | Inf | Inf | NaN | **55** | NaN | NaN | NaN | NaN |
| **6** | NaN | NaN | NaN | NaN | **31** | 14.36 | NaN | 16.44 | NaN | **56** | 51.03 | 41.37 | 32.39 | 73.72 |
| **7** | NaN | NaN | NaN | NaN | **32** | NaN | NaN | NaN | 115.84 | **57** | 44.97 | 21.61 | 44.19 | 63.28 |
| **8** | 1.32 | Inf | 3.21 | 5.03 | **33** | NaN | NaN | NaN | NaN | **58** | Inf | Inf | 283.34 | NaN |
| **9** | 160.41 | 95.64 | 127.82 | NaN | **34** | 67.78 | 30.11 | 59.80 | NaN | **59** | 69.83 | Inf | 62.16 | 80.18 |
| **10** | 32.32 | 21.63 | 29.66 | 41.77 | **35** | NaN | NaN | NaN | NaN | **60** | NaN | NaN | NaN | NaN |
| **11** | NaN | NaN | NaN | NaN | **36** | NaN | NaN | NaN | NaN | **61** | 48.73 | 30.09 | 46.11 | 61.10 |
| **12** | 12.30 | 6.03 | 18.06 | 21.64 | **37** | NaN | NaN | NaN | NaN | **62** | NaN | NaN | NaN | NaN |
| **13** | NaN | NaN | NaN | NaN | **38** | 20.05 | Inf | 15.68 | 152.21 | **63** | 160.32 | 105.20 | 172.29 | NaN |
| **14** | NaN | NaN | NaN | NaN | **39** | NaN | NaN | NaN | NaN | **64** | NaN | NaN | NaN | NaN |
| **15** | 126.20 | 81.50 | 121.26 | 213.96 | **40** | 25.41 | 11.14 | 19.53 | NaN | **65** | NaN | NaN | NaN | NaN |
| **16** | NaN | NaN | NaN | NaN | **41** | NaN | NaN | NaN | NaN | **66** | 268.64 | 134.71 | 278.81 | 242.52 |
| **17** | 5.77 | 4.44 | 8.33 | 5.89 | **42** | 85.63 | 49.99 | NaN | NaN | **67** | 110.35 | 52.11 | 97.86 | 98.60 |
| **18** | 18.31 | 12.81 | 16.62 | 17.36 | **43** | NaN | NaN | NaN | 101.52 | **68** | 351.24 | 73.12 | NaN | NaN |
| **19** | NaN | NaN | NaN | NaN | **44** | 39.34 | NaN | 57.81 | NaN | **69** | 109.77 | NaN | 146.85 | NaN |

| | | | | | | | | | | | | | | |
|---|---|---|---|---|---|---|---|---|---|---|---|---|---|---|
| **20** | NaN | NaN | NaN | NaN | **45** | NaN | NaN | NaN | NaN | **70** | 108.32 | 68.41 | 97.45 | NaN |
| **21** | NaN | NaN | NaN | NaN | **46** | NaN | NaN | NaN | 70.19 | **71** | 125.80 | 59.31 | 105.32 | 99.14 |
| **22** | 17.05 | 5.08 | 5.90 | 27.42 | **47** | NaN | NaN | NaN | NaN | **72** | 121.77 | NaN | 111.35 | NaN |
| **23** | NaN | NaN | NaN | NaN | **48** | 46.12 | 20.20 | 39.58 | NaN | **73** | 239.62 | 65.08 | 100.33 | NaN |
| **24** | NaN | NaN | NaN | NaN | **49** | 25.90 | NaN | 21.63 | NaN | **74** | 22.27 | 5.23 | 17.98 | 20.10 |
| **25** | NaN | NaN | NaN | NaN | **50** | NaN | NaN | NaN | 106.03 | **75** | NaN | NaN | NaN | NaN |
| **76** | NaN | NaN | NaN | NaN | **101** | 137.82 | NaN | 142.89 | NaN | **126** | NaN | NaN | NaN | NaN |
| **77** | 120.25 | 75.03 | 118.19 | 115.84 | **102** | NaN | NaN | NaN | NaN | **127** | 28.80 | Inf | 46.62 | 196.07 |
| **78** | 86.03 | 81.76 | 76.18 | NaN | **103** | 246.97 | 183.80 | 534.47 | 488.71 | **128** | NaN | NaN | NaN | NaN |
| **79** | NaN | NaN | NaN | NaN | **104** | NaN | NaN | NaN | NaN | **129** | Inf | Inf | Inf | Inf |
| **80** | NaN | NaN | NaN | NaN | **105** | NaN | NaN | NaN | NaN | **130** | Inf | Inf | Inf | Inf |
| **81** | NaN | NaN | NaN | NaN | **106** | 46.19 | NaN | 28.53 | NaN | **131** | 292.28 | 91.79 | 345.24 | 444.16 |
| **82** | NaN | NaN | NaN | NaN | **107** | 760.39 | NaN | 593.49 | 730.44 | **132** | 56.18 | NaN | 60.65 | NaN |
| **83** | 140.63 | 81.87 | 120.33 | 152.21 | **108** | NaN | NaN | NaN | NaN | **133** | NaN | NaN | NaN | NaN |
| **84** | 36.79 | NaN | 24.96 | NaN | **109** | 1218.60 | Inf | 2225.77 | Inf | **134** | NaN | NaN | NaN | NaN |
| **85** | NaN | NaN | NaN | NaN | **110** | Inf | Inf | Inf | Inf | **135** | NaN | NaN | NaN | NaN |
| **86** | 90.41 | NaN | 148.37 | NaN | **111** | 87.92 | NaN | 60.68 | 108.57 | **136** | Inf | Inf | Inf | Inf |
| **87** | NaN | NaN | NaN | NaN | **112** | 116.86 | 66.62 | 111.46 | 196.61 | **137** | 47.05 | Inf | 36.66 | 85.72 |
| **88** | 93.63 | 55.30 | 83.16 | 101.52 | **113** | Inf | Inf | Inf | Inf | **138** | NaN | NaN | NaN | NaN |
| **89** | 122.81 | 69.79 | 115.18 | NaN | **114** | NaN | NaN | NaN | NaN | **139** | 378.32 | Inf | 368.29 | Inf |
| **90** | NaN | NaN | NaN | NaN | **115** | Inf | Inf | Inf | Inf | **140** | Inf | Inf | Inf | Inf |
| **91** | 76.17 | 39.39 | 67.33 | 70.19 | **116** | 135.37 | 80.33 | 146.28 | NaN | | | | | |
| **92** | 104.88 | 54.10 | 94.01 | NaN | **117** | NaN | NaN | NaN | NaN | | | | | |
| **93** | 193.11 | 77.53 | 163.64 | NaN | **118** | Inf | Inf | 751.68 | Inf | | | | | |
| **94** | NaN | NaN | NaN | NaN | **119** | Inf | Inf | Inf | Inf | | | | | |
| **95** | 120.54 | 62.43 | 102.95 | 106.03 | **120** | NaN | NaN | NaN | NaN | | | | | |
| **96** | 179.31 | 79.05 | 157.20 | 135.70 | **121** | Inf | Inf | Inf | Inf | | | | | |
| **97** | NaN | NaN | NaN | NaN | **122** | Inf | Inf | Inf | Inf | | | | | |
| **98** | NaN | NaN | NaN | NaN | **123** | NaN | NaN | NaN | NaN | | | | | |

| | | | | | | | | | |
|---|---|---|---|---|---|---|---|---|---|
| **99** | 75.99 | 51.83 | 71.47 | 98.03 | **124** | Inf | NaN | Inf | Inf |
| **100** | 149.96 | 102.31 | 137.96 | 160.21 | **125** | Inf | Inf | Inf | Inf |

**Table S3:** Residue specific a$K_D$ values plotted in heat map in Figure 5B.
